# Lattice defects induced by microtubule-stabilizing agents exert a long-range effect on microtubule growth by promoting catastrophes

**DOI:** 10.1101/2021.02.11.430743

**Authors:** Ankit Rai, Tianyang Liu, Eugene A. Katrukha, Juan Estévez-Gallego, Ian Paterson, J. Fernando Díaz, Lukas C. Kapitein, Carolyn A. Moores, Anna Akhmanova

## Abstract

Microtubules are dynamic cytoskeletal polymers that spontaneously switch between phases of growth and shrinkage. The probability of transitioning from growth to shrinkage, termed catastrophe, increases with microtubule age, but the underlying mechanisms are poorly understood. Here, we set out to test whether microtubule lattice defects formed during polymerization can affect growth at the plus end. To generate microtubules with lattice defects, we used microtubule-stabilizing agents that promote formation of polymers with different protofilament numbers. By employing different agents during nucleation of stable microtubule seeds and subsequent polymerization phase, we could reproducibly induce switches in protofilament number and induce stable lattice defects. Such drug-induced defects led to frequent catastrophes, which were not observed when microtubules were grown in the same conditions but without a protofilament number mismatch. Microtubule severing at the site of the defect was sufficient to suppress catastrophes. We conclude that structural defects within microtubule lattice can exert effects that can propagate over long distances and affect the dynamic state of the microtubule end.

## Introduction

Microtubules are cytoskeletal polymers that rapidly switch between phases of growth and shortening, and this behavior, termed dynamic instability, plays a crucial role in the formation, maintenance and reorganization of microtubule arrays during cell division, migration and differentiation (1, 2). The transition from growth to shrinkage, an event called catastrophe, is known to occur when the protective cap of GTP-bound tubulin subunits is reduced or lost, but the underlying mechanisms are still subject of investigation (3, 4). One interesting property of microtubules is that the frequency of catastrophes depends on microtubule age: microtubules that are growing for a longer time have a higher chance to switch to depolymerization (5, 6). Changes occurring at the microtubule end, such as loss of individual protofilaments or end tapering have been shown to promote catastrophe (7–9). In principle, it is also possible that the catastrophe frequency at the plus end is affected by structural features in the microtubule lattice further away from the tip, but this possibility has so far remained untested.

Structural studies have established that tubulin can form tubes with different protofilament numbers, dependent on the species, nucleation template, presence of different microtubule-associated proteins and other properties of the polymerization reaction (e.g. (10), reviewed in (11)). An important consequence of the structural plasticity of the microtubule lattice (12) is the formation of lattice defects, such as sites where a microtubule gains or loses one or more protofilaments (10, 13–16). A recent cryo-electron tomography analysis showed that in some cell types, such as *Drosophila* neurons, variations and transitions in protofilament number are readily detectable (17) and are thus likely to be physiologically relevant. Switches in protofilament number can be introduced during microtubule growth, and their presence may affect microtubule dynamics in different ways. For example, defects can be repaired through tubulin incorporation, and the resulting islands of GTP-tubulin can trigger microtubule rescue (18–21). On the other hand, the presence of defects could potentially also induce catastrophes (as proposed in ref (13)), since conformational properties of the microtubule lattice might propagate over some distances (22).

To study the relationship between lattice defects and microtubule catastrophes, one should be able to directly correlate the presence of defects with the dynamics of microtubule ends. We recently found that fluorescent analogues of microtubule-stabilizing agents (MSAs) can be used to induce microtubule lattice defects that can be visualized by fluorescence microscopy. When present at low concentrations, MSAs preferentially bind to microtubule plus ends that enter a “pre-catastrophe” state (23), which is manifested by the gradual loss of the GTP cap and reduced recruitment of End Binding (EB) proteins that detect GTP-bound microtubule lattice (24–27). Strong accumulation of MSAs at pre-catastrophe microtubule ends leads to the formation of stabilized patches of microtubule lattice, where the tube is incomplete and keeps incorporating GTP-tubulin but is not fully repaired (23). When microtubules switch to depolymerization, such persistent lattice defects, which coincide with the hotspots of MSA binding, can induce repeated rescues and, therefore, they were termed “stable rescue sites” (23).

Here, we used MSA-induced lattice defects to address two questions. First, what prevents complete repair of an MSA-induced persistent lattice defect? And second, does the presence of such a persistent defect affect the dynamics of the microtubule plus end? Since different MSAs are known to affect the number of protofilaments (14, 28–34), we hypothesized that persistent lattice defects could be associated with the changes in protofilament number and thus could not be fully repaired for geometrical reasons. We tested this idea by generating stable microtubule seeds with one MSA and then elongating them in the presence of another MSA, with the same or different preference for protofilament number. Use of fluorescent MSAs allowed us to directly follow drug binding. We found that pre-catastrophe microtubule ends accumulated MSAs in all conditions; however, the outcome of drug binding was different. When there was no mismatch in protofilament number between the seeds and the elongation conditions, drug accumulations were short in duration and length, and microtubule growth beyond such sites was processive. In contrast, when, based on the MSA properties, a mismatch in protofilament number could be expected, large and persistent drug accumulations were formed. The existence of such mismatches was confirmed by cryo-electron microscopy (cryo-EM) and by measuring microtubule growth rate, which became higher with increasing protofilament number. When microtubule ends extended beyond a mismatch-containing lattice defect, they displayed elevated catastrophe frequency. Laser-mediated severing of a microtubule at the site of the persistent defect reduced catastrophe frequency at the plus end. Our data demonstrate that local perturbations in microtubule structure can affect the state of the dynamic end at a distance of several micrometers.

## Results

### Protofilament number affects microtubule growth rate

We first set out to test whether microtubules with different protofilament numbers display different dynamic properties. Protofilament number changes in response to the nucleotide bound to the regulatory E-site or the presence of MSAs (14, 28–34). Therefore, to generate microtubules with different protofilament numbers, we prepared microtubule seeds with the slowly hydrolysable GTP analog GMPCPP or with different microtubule-stabilizing drugs. Using X-ray fiber diffraction and cryo-EM, we confirmed previous observations showing that microtubule seeds generated in the presence of Taxol have predominantly 12/13 protofilaments (pf), whereas 14pf microtubules were observed in the presence of GMPCPP and Docetaxel (Fig. 1A, B) (14, 31–34). Microtubules stabilized with Alexa_488_-Epothilone B also had 14pf, whereas protofilament number shifted towards 15pf in the presence of Discodermolide and 15/16pf with Fchitax-3 (Fig. 1A, B). We next used these different stabilized microtubule seeds to grow dynamic microtubules and observed their behavior using Total Internal Reflection Fluorescence Microscopy (TIRFM) as described previously (35, 36) (Fig. S1A). Assays were performed either with tubulin alone or with the addition of mCherry-EB3, which serves as a microtubule plus end marker and increases both the growth rate and catastrophe frequency in assays with purified tubulin (37) (Fig. 1C, Fig. S1B). In these assays, MSAs were used to prepare stable seeds but were not added during polymerization. Both with and without EB3, we found that microtubule growth rate increased with the protofilament number (Fig. 1D upper panel, Fig. S1C). Calculation of the critical concentration *C_c_* of microtubule polymerization based on fitting of the dependence of the growth rate on tubulin concentration (38) indicated that *C_c_* is higher for microtubules with fewer protofilaments (Fig. 1E). Catastrophe frequency showed some variability between different conditions but was independent of protofilament number (Fig. 1D lower panel, Fig. S1D). As described previously (36), very few rescues were seen when microtubules were grown from GMPCPP seeds, while occasional rescues were found when drug-stabilized seeds were used, likely because some drug molecules could diffuse from the seeds and bind to the dynamic lattice.

**Fig. 1.**
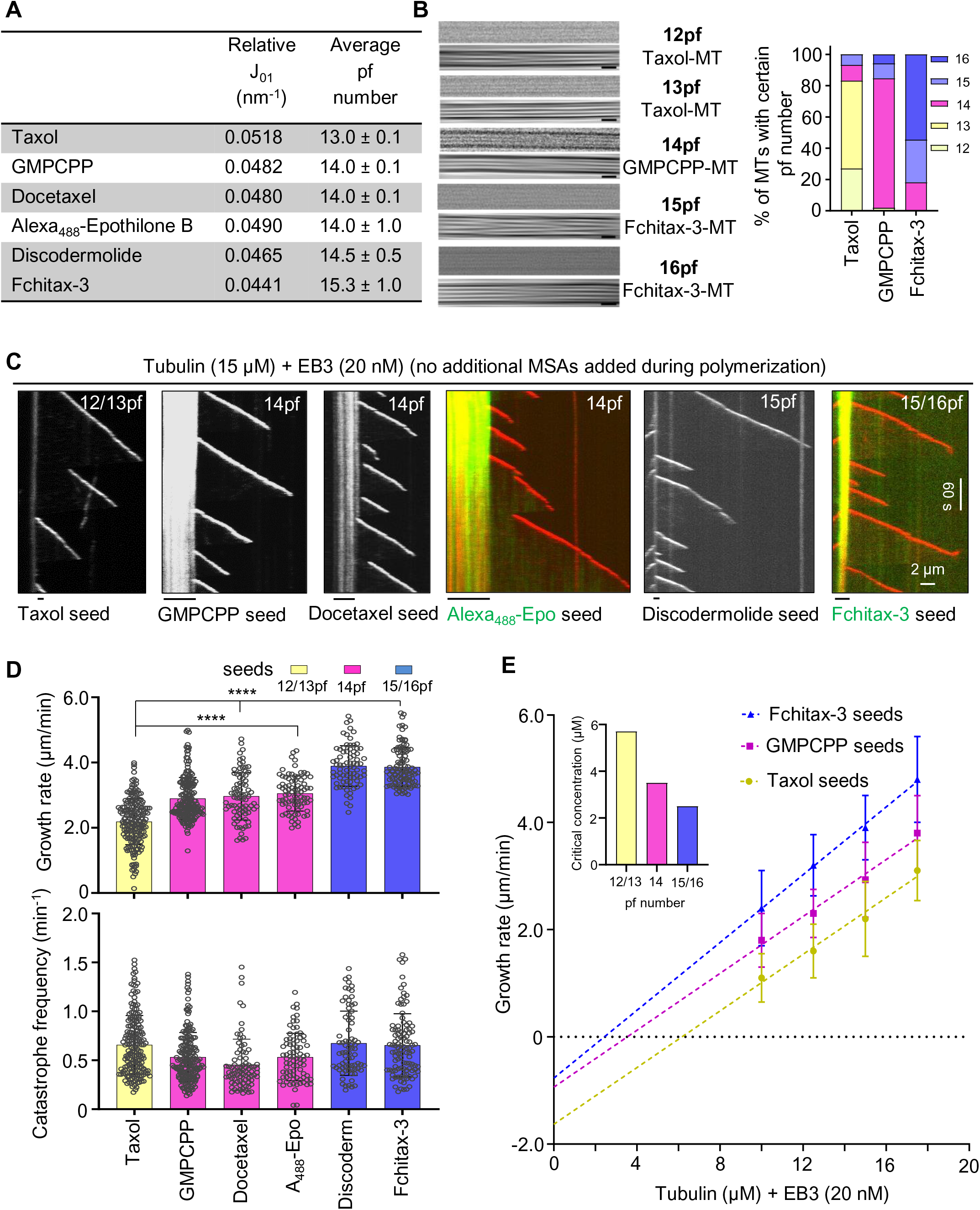
Microtubule protofilament number affects microtubule growth rate. **A)** X-ray fiber diffraction measurements for protofilament (pf) number quantification of microtubules polymerized in the presence of GMPCPP or different MSAs. For each condition, a total of 24 diffraction images were averaged and background subtracted using ImageJ software. **B)** Representative raw cryo-EM images (in the presence of Taxol, GMPCPP and Fchitax-3) and their filtered versions emphasizing their Moiré patterns. Scale bar, 25 nm. Bar graph shows microtubule protofilament number distribution for different conditions determined by Moiré pattern visualization for each microtubule. Microtubule population: n = 89 for Taxol; n = 52 for GMPCPP; n = 77 for Fchitax-3. **C)** Representative kymographs showing microtubule dynamics in the presence of seeds (stabilized with GMPCPP or indicated MSAs) with different protofilament numbers, supplemented with tubulin (15 μM) and mCherry-EB3 (20 nM) in the absence of any additional MSAs added during the reaction. **D)** Quantification of growth rates (upper panel) and catastrophe frequencies (lower panel) in the presence of seeds with different protofilament numbers as presented in panel C. From left to right, n = 193, 196, 82, 81, 76, 104 growth events. N = 2 independent experiments for Docetaxel, Alexa_488_-Epothilone B and Discodermolide, N = 3 independent experiments for Taxol, GMPCPP and Fchitax-3. Error bars represent SD; ****, p <0.0001, Mann-Whitney U test. **E)** Microtubule growth rate as a function of tubulin concentration (10 – 17.5 μM) from seeds with different protofilament numbers. Error bars represent SD, Cc is the critical concentration (shown in inset) calculated based on the linear fits of the data, N = 3 independent experiments. Microtubule growth events; n = 280, 161 and 221 for 10 μM, n = 369, 254, 193 for 12.5 μM, n = 193, 196, 104 for 12.5 μM, n = 243, 208 and 214 for 17.5 μM for Taxol, GMPCPP and Fchitax-3, respectively.

Importantly, in these conditions, the nature of the drug present in the seeds had no effect on the polymerization rate - for example, all microtubules grown from 14pf seeds, including GMPCPP-stabilized ones, polymerized with the same speed. To further exclude that the observed differences in growth rates were caused by the drugs diffusing from the seeds, we labeled the seeds with different protofilament numbers in different colors and grew microtubules from two types of seeds on the same coverslip (Fig. S2A). In these experiments, two types of seeds were exposed to exactly the same reaction mix, including the drugs that might be present in solution. Importantly, we observed that microtubules still displayed growth rates characteristic for their protofilament number (Fig. S2A). For example, in the same reaction mix, microtubules grew from Taxol-stabilized seeds (12/13pf) slower than from GMPCPP (14pf) seeds (Fig. S2A). We also investigated whether microtubule depolymerization rate depended on protofilament number but found no clear correlation: we observed that microtubules grown from 14pf or 15/16pf seeds depolymerized at the same rate, but slower than microtubules grown from Taxol-stabilized (12/13pf) seeds (Fig. S2B). We conclude that when the growth conditions are the same, microtubule polymerization rate can be used to infer protofilament number.

### Effects of MSAs on microtubule dynamics depend on the protofilament number in the seeds

Next, we investigated how microtubule dynamics would be affected by adding during polymerization an MSA with a protofilament number preference that was either the same (matching conditions) or different (mismatching conditions) than the one used during seed preparation (Fig. 2A). Drug concentrations in the range 50-400 nM were used, because higher concentrations induced spontaneous microtubule nucleation, and we thus could not ensure that all observed microtubules grew from the pre-existing seeds with the known protofilament number. We observed three types of dynamic microtubule behaviors. The first type of dynamics was semi-processive growth interrupted by short (0.2 - 0.5 µm) depolymerization events followed by rapid rescues (“semi-processive growth”, Fig. 2B, C, Fig. S3A, C). The second type of dynamic behavior was manifested by frequent catastrophes followed by long (>0.5 µm) depolymerization events and repeated rescues at the same site (stable rescue site, which will be termed here “SRS dynamics”, Fig. 2B, D, Fig. S3B, C), The third type of dynamic behavior was characterized by catastrophes followed by long (>0.5 µm) depolymerization events and randomly distributed rescues (termed “random rescues”, Fig. 2B, Fig. S4A). At 100 nM MSA concentrations, there was a clear difference in the occurrence of a particular type of dynamics, which depended on the combination of MSAs used for preparing the seeds and their elongation. When the protofilament number was expected to be the same (“matching conditions”, Fig. 2A), semi-processive growth with very short depolymerization events strongly predominated (Fig. 2B, C, Fig. S3A, C, short growth perturbations highlighted by asterisks). In contrast, when the seeds were elongated in the presence of an MSA that had a protofilament number preference that was different from that of the MSA used for seed stabilization (“mismatching conditions”, Fig. 2A), SRS dynamics (highlighted by white dashed lines) or random rescues (highlighted by yellow arrows) with long depolymerization events were observed (Fig. 2B,D, Fig. S3B,C, S4A). When higher MSA concentrations were used during microtubule growth, random rescues were predominantly observed for mismatching conditions (Fig. S4A), likely because rescues became more frequent, and depolymerization events were thus not long enough to reach the preceding stable rescue site. When MSA concentration was reduced, some microtubules displayed random rescues in matching conditions, because depolymerization events became longer but were typically still followed by rapid rescues (Fig. S4B).

**Fig. 2.**
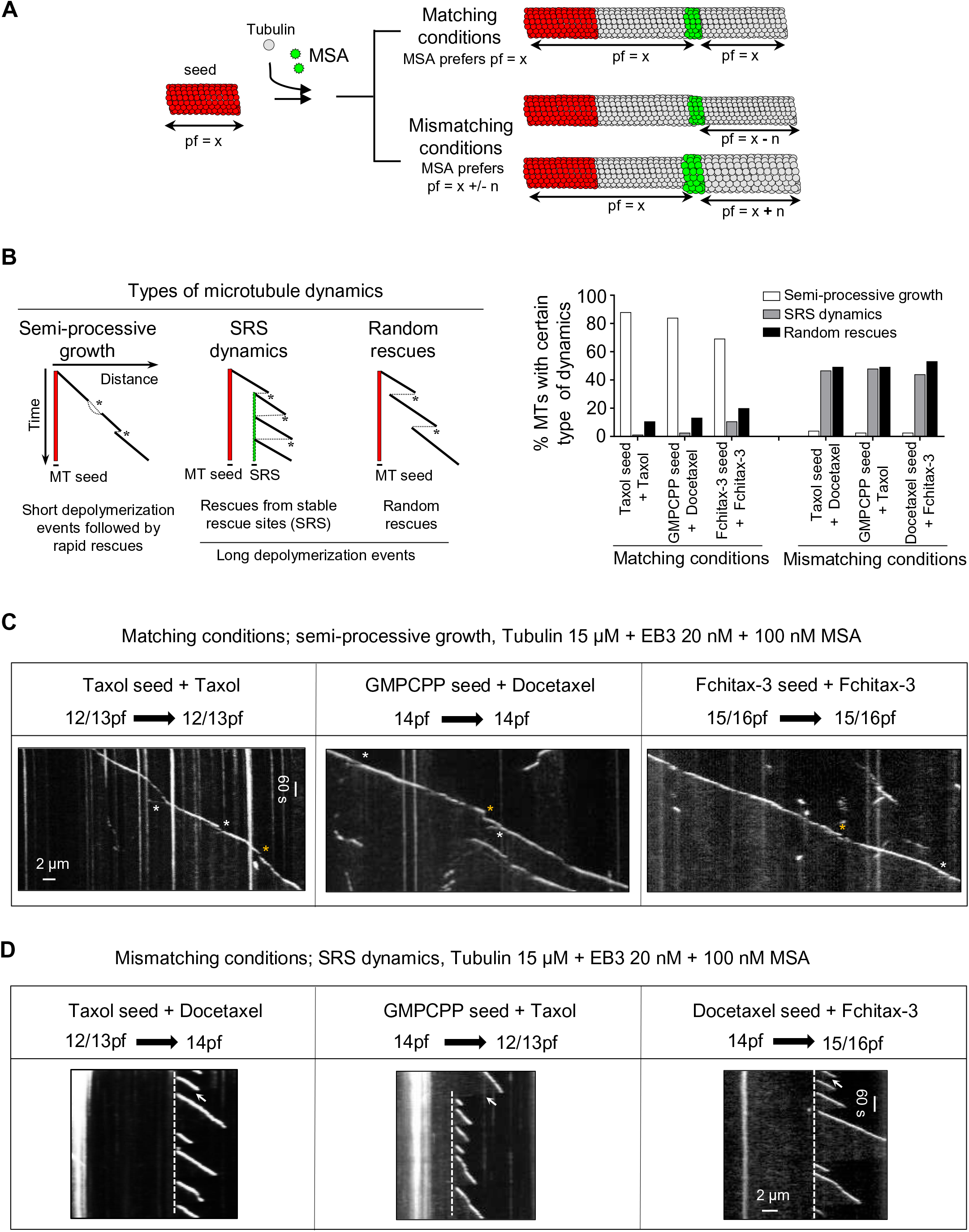
Protofilament number mismatch between the seed and growth conditions affects microtubule dynamics. **A)** A scheme illustrating microtubule growth from seeds in the presence of MSAs with the same or different protofilament number preference. MSA binding zones are shown in green. **B)** (Left) A cartoon depicting kymographs that correspond to three different types of microtubule dynamics observed in different conditions. Microtubule dynamics is defined as SRS dynamics if a microtubule regrows at least 3 times from the same site after undergoing catastrophes. A random rescue is a single rescue event after depolymerization episode that is longer than 0.5 μm. Asterisks highlight catastrophe events in each condition. (Right); Quantification of microtubule dynamics observed in the indicated conditions, as represented in the cartoon and in panels (C and D). n = 75 microtubule seeds in all the conditions, N = 3 independent experiments. **C, D)** Representative kymographs showing microtubule dynamics in the indicated conditions. In matching conditions, short growth perturbation events followed by rapid rescues are highlighted (white asterisks highlight split comets and yellow asterisks highlight depolymerization events with the length 0.2-0.5 μm). Stable rescue sites in mismatching conditions are highlighted by white stippled lines. A white arrow highlights a long depolymerization event (>0.5 μm). N = 3 independent experiments.

Importantly, microtubules grown in the presence of the same MSA showed very different dynamics dependent on the seeds used. For example, microtubules grown in the presence of 100 nM Taxol displayed semi-processive growth when extending from Taxol-stabilized seeds (12/13pf), but SRS dynamics or random rescues when grown from GMPCPP-stabilized (14pf) or Fchitax-3 stabilized (15/16pf) seeds (Fig. 2B-D, Fig. S3A-C). In contrast, in the presence of Docetaxel, microtubules grew semi-processively from GMPCPP or Docetaxel-stabilized (14pf) seeds but showed SRS dynamics or random rescues when grown from Taxol- or Fchitax-3 stabilized seeds (Fig. 2B-D, Fig. S3A-C). Similar results were also obtained when microtubules were grown without mCherry-EB3, although the absence of a plus-end marker made the detection of short depolymerization events less reliable (Fig. S5). For subsequent analyses, we therefore focused on the data obtained with MSA concentrations of 100 nM in the presence of 20 nM mCherry-EB3. Together, these results demonstrate that the (mis)match between the protofilament number of the seed and the number preferred by the MSA present during elongation has a strong effect on microtubule growth dynamics, even when the microtubule tip is far away from the seed.

### Larger and more persistent drug accumulations are observed in mismatching conditions

To better understand the origin of the striking differences in the observed microtubule dynamics in matching and mismatching conditions, we visualized drug binding using fluorescent drug analogues. Stable rescue sites coincide with the formation of drug accumulation hotspots, which initiate directly behind a growing microtubule plus end entering a pre-catastrophe state that can be distinguished by the reduction in EB3 signal (23) (Fig. 3A, B). Microtubule lattice zones with high drug affinity can extend for several micrometers, but then abruptly stop (Fig. 3A, B), and previous work suggested that they might represent incomplete tubes, which stop binding the drug when they close (23).

**Fig. 3.**
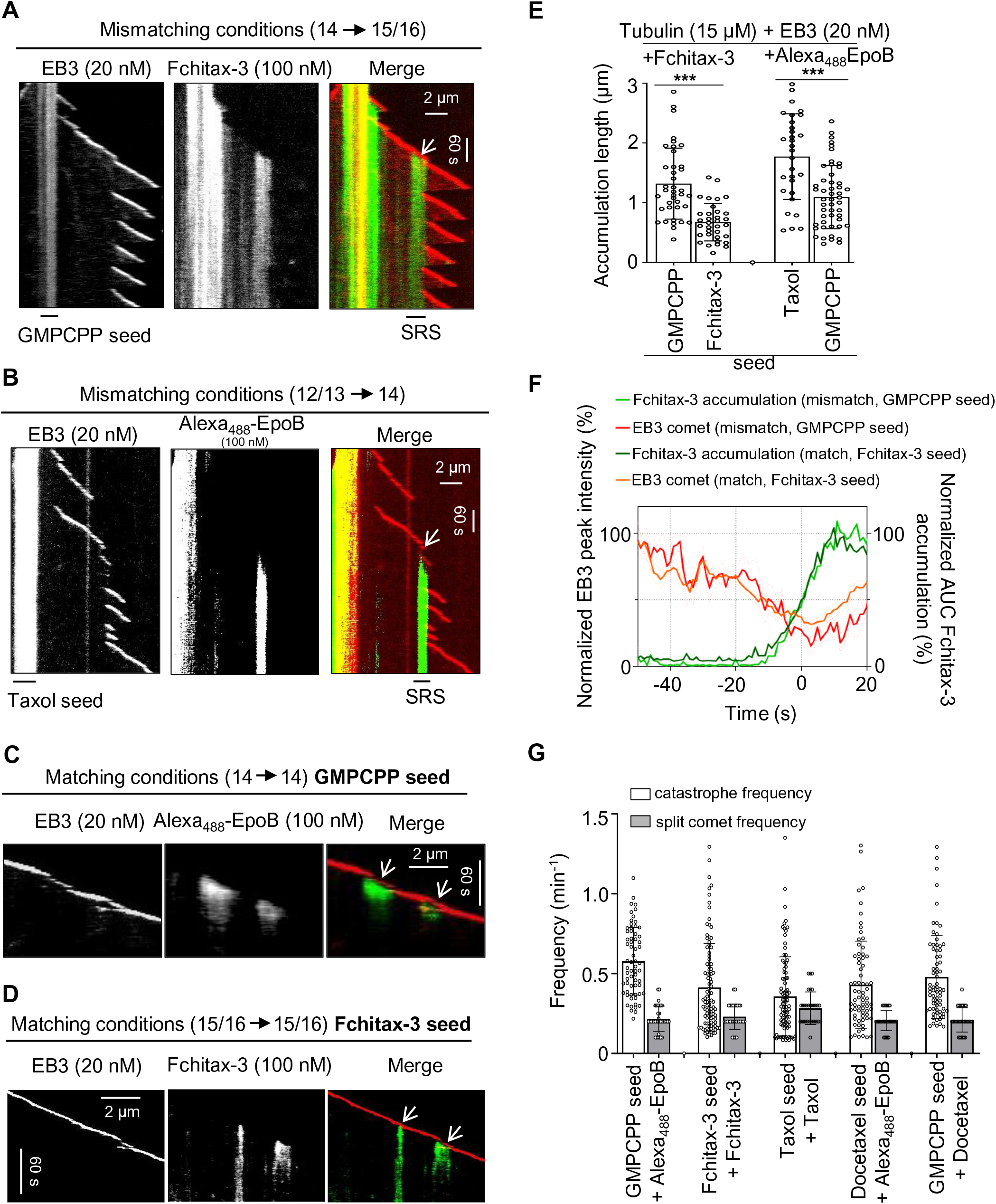
Extent of MSA accumulation at pre-catastrophe microtubule tips depends on the match between the seed and growth conditions. **A, B)** Representative kymographs illustrating drug accumulations in mismatching conditions. Microtubules were grown from GMPCPP seeds (A) or Taxol seeds (B) in the presence of 15 μM tubulin and 20 nM mCherry-EB3 with Fchitax-3 (100 nM) (A) or Alexa_488_-Epothilone B (100 nM) (B). White arrows indicate stable rescue sites with drug accumulations. **C, D)** Representative kymographs illustrating drug accumulations in matching conditions. Microtubules were grown from GMPCPP seeds (C) or Fchitax-3 seeds (D) in the presence of 15 μM tubulin and 20 nM mCherry-EB3 with Alexa_488_-Epothilone B (100 nM) (C) or Fchitax-3 (100 nM) (D). Split comets are indicated with white arrows. **E)** Quantification of drug accumulation lengths in the indicated conditions. Error bars indicate SD; from left to right, n = 39, 34, 30 and 50, N = 3 independent experiments, ***, p <0.001. **F)** Time plot of averaged normalized maximum intensity of fitted EB3 comet (orange, red) and normalized area under the curve of fitted Fchitax-3 (light and dark green) intensity profiles in mismatching conditions (as shown in panel A, n = 9 kymographs from 5 experiments) and in matching conditions (as shown in panel D, n = 38 kymographs from 5 experiments). Individual curves were aligned by maximizing cross-correlation between Fchitax-3 time curves. Error bars represent SEM. **G)** Frequencies of catastrophes (calculated as frequency of all growth perturbations including “split comet” events) and split comet events in different matching conditions. From left to right, n = 65, 100, 91, 71, and 70 for catastrophe frequencies for the indicated conditions and n = 60, 66, 73, 62 and 63 “catching up” events from 30 microtubules for split comet frequencies. N = 3 independent experiments. Error bars represent SD.

In mismatching conditions, such as GMPCPP seeds (14pf) elongated in the presence of Fchitax-3 (15/16pf) or Taxol-stabilized seeds (12/13pf) elongated in the presence of Alexa_488_-Epothilone B (14pf), we observed large and persistent drug accumulation zones that coincided with stable rescue sites (Fig. 3A, B). In contrast, in matching conditions (GMPCPP-stabilized seeds elongated in presence of Alexa_488_-Epothilone B (14pf) or Fchitax-3-stabilized seeds elongated in the presence of Fchitax-3 (15/16pf), drug accumulation patches were short in duration and length (Fig. 3C-E, Fig. S6A). In both conditions, drug binding always initiated directly behind a microtubule plus end after it started to lose mCherry-EB3 signal, indicating a growth perturbation. The kinetics of the reduction of EB3 signal before drug binding showed considerable variability but was similar in matching and mismatching conditions (Fig. 3F), indicating that in both situations, microtubules could enter a pre-catastrophe state. The initial phase of drug accumulation was also very similar for matching and mismatching conditions (Fig. 3F).

Importantly, growth perturbations in matching conditions were typically of limited duration and were often accompanied by the emergence of a second, faster comet at the rear of the drug accumulation site (Fig. 3C, D, Fig. S6A). Previous work showed that a “catching up” comet appears when some protofilaments at the growing microtubule tip are stalled whereas the others keep elongating. When the stalled protofilaments start to regrow, a faster rear comet emerges and ultimately fuses with the leading one (8, 39). We observed split comets during growth perturbations in all tested matching conditions, also when non-fluorescent MSAs were used (Fig. S6B, C). Clear split comets were seen in 38-79% of all catastrophe events detected in matching conditions (Fig. 3G, all growth perturbations with the length >0.2 µm, including “split comet” events, were considered as catastrophes as indicated by asterisks in Fig. 2B). Since the two comets must be located at a significant distance from each other to be registered as a “split comet” by fluorescence microscopy, these numbers are likely underestimates of the actual frequency of such events. We conclude that in matching conditions, growth perturbations are followed by the formation of “catching up” comets, which likely help to restore a normally growing microtubule plus end, leading to semi-processive microtubule growth.

In mismatching conditions, drug accumulation zones were longer and much more persistent (Fig. 3A-E). The binding density of Fchitax-3 in mismatching conditions (∼1-2 drug molecules per 8 nm (ref (23)) was higher than in matching conditions (0.3-0.9 molecules per 8 nm microtubule length, Fig. S6D). This can be explained by the fact that in mismatching conditions, drug accumulations expand for a longer time, possibly because a normal tube is more difficult to restore and protofilaments continue growing as a sheet or some other microtubule end structure that promotes drug binding. Thus, a mismatch between the protofilament number preference of the MSA used to prepare the seeds and to elongate them inhibits the restoration of a growing microtubule plus end after a growth perturbation has occurred and a drug accumulation has formed.

### Lattice defects observed in mismatching conditions are associated with switching of protofilament number

We hypothesized that binding of the drug to a pre-catastrophe microtubule tip either induces or stabilizes tubes with a protofilament number fitting with the specific preference of the drug used. In mismatching conditions, this would cause a protofilament number switch occurring at the stable rescue site, and this could explain why such sites do not get fully repaired. If this hypothesis is correct, a microtubule will be expected to grow with the speed characteristic for the number of protofilaments present in the seed before the stable rescue site, but with a speed characteristic for the MSA present in the growth reaction after it. We found that in matching conditions, the addition of any MSA caused an increase in microtubule growth rate, but the correlation between the rate and protofilament number was retained (Fig. 4A, upper panel). Interestingly, microtubules with SRS dynamics in mismatching conditions displayed growth rates characteristic for the seed before the rescue site and the growth rate matching better that of the MSA used in the growth reaction after the stable rescue site. For example, when GMPCPP seeds (14pf) were elongated in the presence of Fchitax-3 (15/16pf) or Taxol (12/13pf), the growth rate was characteristic for 14pf microtubules before the stable rescue site but was elevated after a stable rescue site in the presence of Fchitax-3 and decreased in the presence of Taxol (Fig. 4A, upper panel). In contrast, in matching conditions, growth rate before and after a growth perturbation did not change (Fig. S7A). It should be noted, however, that the changes in growth rate did not completely match the speeds of microtubule growth when the same MSA was used for seed stabilization and elongation (Fig. 4A, upper panel). This likely reflects variability in protofilament number after the switch at a stable rescue site.

**Fig. 4.**
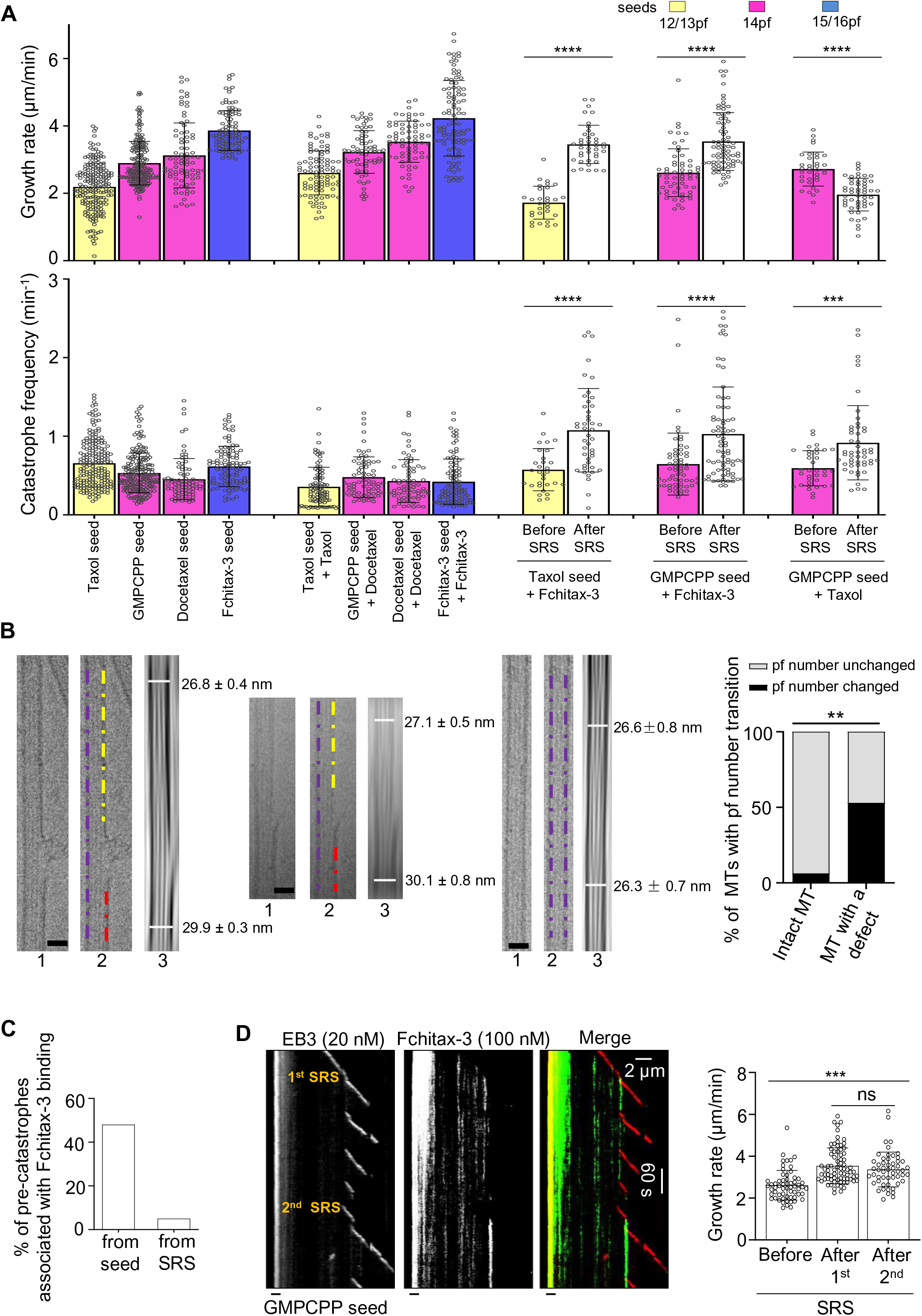
Lattice defects observed in mismatching conditions are associated with switching of protofilament number. **A)** Quantification of growth rates (upper panel) and catastrophe frequencies (lower panel, calculated as frequency of all growth perturbations including “catching up” events) for the indicated conditions. For better comparison, data from Figure 1D are presented again in the present panels. n = 193, 196, 82, 104 microtubule growth events for microtubules polymerized from Taxol, GMPCPP, Docetaxel, Fchitax-3 seeds. n = 91, 70, 71, 100 in the presence of Taxol seed + Taxol (100 nM), GMPCPP seed + Docetaxel (100 nM), Docetaxel seed + Docetaxel (100 nM), Fchitax-3 seed + Fchitax-3 (100 nM). n = 30 and 40 for Taxol seed + Fchitax-3 (100 nM) before and after stable rescue site (SRS) formation. n = 60 and 75 for GMPCPP seed + Fchitax-3 (100 nM) before and after SRS formation. n = 35 and 51 for GMPCPP seed + Taxol (100 nM) before and after SRS formation. N = 3 independent experiments. Error bars represent SD. The colors of the bars and pf values indicate the protofilament number preferences of a particular drug used. ***, p <0.001, ****, <0.0001, Mann-Whitney U test. **B)** Representative raw cryo-EM images and their filtered versions with enhanced microtubule Moiré patterns showing the diameter difference (pf number transition) for microtubules grown from GMPCPP seeds in the presence of Fchitax-3 (mismatching condition, Figure 3A). The left and central sets of panels show defect-containing microtubules and the right one shows a microtubule with no visible defects. In each panel, Image 1 is the raw cryo-EM image of one microtubule. Scale bar, 25 nm. Image 2 is the same cryo-EM image with purple, yellow and red dash lines highlighting diameter difference on either side of a defect. Image 3 shows the diameters for two sides of microtubule. Diameters were measured in Fiji. Percentage of microtubule with protofilament number transition in microtubules with no visible defects (n = 19) and microtubules with sheet-like defects (n = 17). The percentage differences were evaluated by two-sided Fisher’s exact test. **, p value = 0.0023. **C)** Bar graph shows quantification of frequency of drug accumulations at pre-catastrophe microtubule ends (identified by the strong reduction or complete loss of EB3 intensity, as shown in Figure 3F) growing either from a seed or after a stable rescue site (SRS). **D)** Representative kymographs showing occurrence of a second stable rescue site (SRS) after the formation of the first SRS within the same microtubule. The experiment was performed with GMPCPP seeds in the presence of 15 μM tubulin, 20 nM EB3 and 100 nM Fchitax-3. Bar graph shows quantification of microtubule growth rates before (n = 60) and after the first (n = 75) and the second (n = 53) stable rescue site. N = 3 independent experiments, ns = not significant, ***, p <0.001, Mann-Whitney U test.

Further support for the occurrence of protofilament number switching at the stable rescue sites was obtained by cryo-EM. In our previous study (23), we found that microtubule lattice discontinuities corresponding to Fchitax-3 accumulations can be detected by cryo-EM. Here, we analyzed these data focusing on the microtubule diameter at the two sides of a lattice defect and found that protofilament number changed in ∼53% of such cases, whereas in microtubules lacking defects switches in protofilament number were rare (Fig. 4B, Fig. S7B). Since not all defects detected by cryo-EM might represent stable rescue sites observed by fluorescence microscopy, this number likely represents an underestimate of the actual switching of protofilament number at stable rescue sites. Thus, MSAs present during microtubule elongation can induce a switch to their preferred protofilament number at the stable rescue site.

### Protofilament number mismatch between the seed and the growing microtubule lattice promotes catastrophes

Having established that stable rescue sites can correspond to regions where a switch in protofilament number takes place, we next asked how such sites affect growth of microtubules extending beyond them. Interestingly, we found that whereas catastrophe frequency (calculated as frequency of all growth perturbations >0.2 µm including “catching up” events) remained constant for microtubules growing in matching conditions, it was strongly increased for the growth events occurring after the stable rescue site (Fig. 4A, lower panel). The increase in catastrophe frequency was similar for microtubules that switched to higher (e.g. from 12/13pf or 14pf to 15/16pf) or lower (from 14pf to 12/13pf) number of protofilaments (Fig. 4A, lower panel). This observation was surprising, because one would expect that after switching to the protofilament number “preferred” by the MSA present in solution, a microtubule will be further growing in matching conditions and should thus display semi-processive growth without long depolymerization events. However, this was not the case: microtubule plus ends entering a pre-catastrophe state after a stable rescue site typically did not accumulate MSAs and simply switched to depolymerization, which proceeded all the way back to the preceding stable rescue site (Fig. 3A, B). Whereas 48% of all catastrophe and pre-catastrophe events (distinguished by strong reduction or complete loss of the EB3 signal) occurring during microtubule outgrowth from the seed led to drug accumulation and formation of a stable rescue site, only 5% of such events occurring after a stable rescue site triggered drug accumulation and microtubule stabilization (Fig. 4C). Formation of a secondary stable rescue site was thus quite rare: for example, when microtubules extended from GMPCPP-stabilized seeds in the presence of Fchitax-3, the formation of secondary stable rescue site was seen only in 11% of all observed microtubules (Fig. 4D). This suggests that some properties of a pre-catastrophe microtubule plus end extending after a stable rescue site (after a microtubule has incorporated a lattice defect) are different from those of microtubule ends growing directly from the seed.

To explore the underlying mechanism, we analyzed the intensities of EB3 comets and microtubule tip tapering during growth episodes from the GMPCPP-stabilized seed (14pf) either in the absence of drugs (control) or after a stable rescue site induced by Discodermolide (14/15pf, mismatching conditions). We note that Fchitax-3 or Alexa_488_-Epothilone B could not be used in these experiments because we used green (HiLyte 488^TM^ labeled tubulin) fluorescence to obtain tubulin profiles. We focused on the 40 s time interval preceding a catastrophe in order to determine whether the pre-catastrophe plus ends of microtubules growing directly from the seed or from a stable rescue site are somehow different. Microtubule end tapering was similar in both conditions (Fig. S7C). The reduction of EB3 signal during catastrophe onset, which is expected to reflect the kinetics of GTP cap loss during catastrophe initiation, was also quite similar, although the curve was somewhat less steep for microtubules growing from seeds compared to microtubules elongating beyond a stable rescue site (Fig. S7D). A higher-resolution analysis would be needed to determine what makes pre-catastrophe microtubule tips growing from seed different from those elongating after a stable rescue site.

### Microtubule severing at the lattice defect site suppresses catastrophes

Why do microtubules growing beyond a stable rescue site display more catastrophes? One possibility is that a lattice defect with a different number of protofilaments on the two sides affects the growth at the plus end through a long-range conformational alteration or mechanical strain. If this is the case, severing the microtubule at the site of the defect should reduce catastrophe frequency. To test this idea, we performed microtubule severing on a TIRF microscope using a pulsed 532 nm laser and observed the dynamics of the severed part of the microtubule (Fig. S8A). The severed microtubule fragment was no longer attached to the coverslip. However, due to the presence of methylcellulose, which increases the viscosity of the solution and dampens fluctuations, most severed microtubule fragments did not float away, but stayed close to the surface and could still be observed by TIRF microscopy (Figure S8B, Video S1). In some cases, they underwent diffusive movements, however, movements of the whole microtubule segment could be easily distinguished from microtubule growth and shortening by the synchronous displacement of fluorescent speckles present along the microtubule shaft (Fig. S8B). After microtubule severing in the absence of MSAs (microtubules polymerized in the presence of GMPCPP seeds with 15 µM tubulin and 20 nM EB3), freshly generated microtubule plus ends typically depolymerized, whereas freshly generated minus ends displayed heterogeneous behavior. The poor survival of microtubules after severing and the heterogeneity in minus-end dynamics precluded a meaningful analysis of the severing data in the absence of MSAs.

In the presence of an MSA such as Fchitax-3, the severed microtubule ends were typically quite stable. To generate microtubules with defects that would be visible by fluorescence microscopy, we grew microtubules in mismatching conditions from GMPCPP-stabilized seeds (14pf) in the presence of 100 nM Fchitax-3 (15/16pf), rhodamine-labeled tubulin and 20 nM mCherry-EB3. After photoablation, the newly generated plus end, which remained attached to the seed, as well as the new minus end and the pre-existing plus end of the microtubule fragment that was detached from the seed could all polymerize (Fig. 5A). Imaging before photoablation and the photoablation itself induced significant photobleaching, and the tubulin freshly incorporated at the growing microtubule ends after severing could be readily observed because its signal was brighter; growing plus ends were additionally visualized by the accumulation of mCherry-EB3, that was detected in the same fluorescent channel as tubulin (Fig. 5A). Even when the severed end underwent some displacement after photoablation, the plus- and the minus end could be easily distinguished from each other by their growth rate, and the remnant of the drug accumulation zone at the stable rescue site was also visible (labeled as SRS in Fig. 5A).

**Fig. 5.**
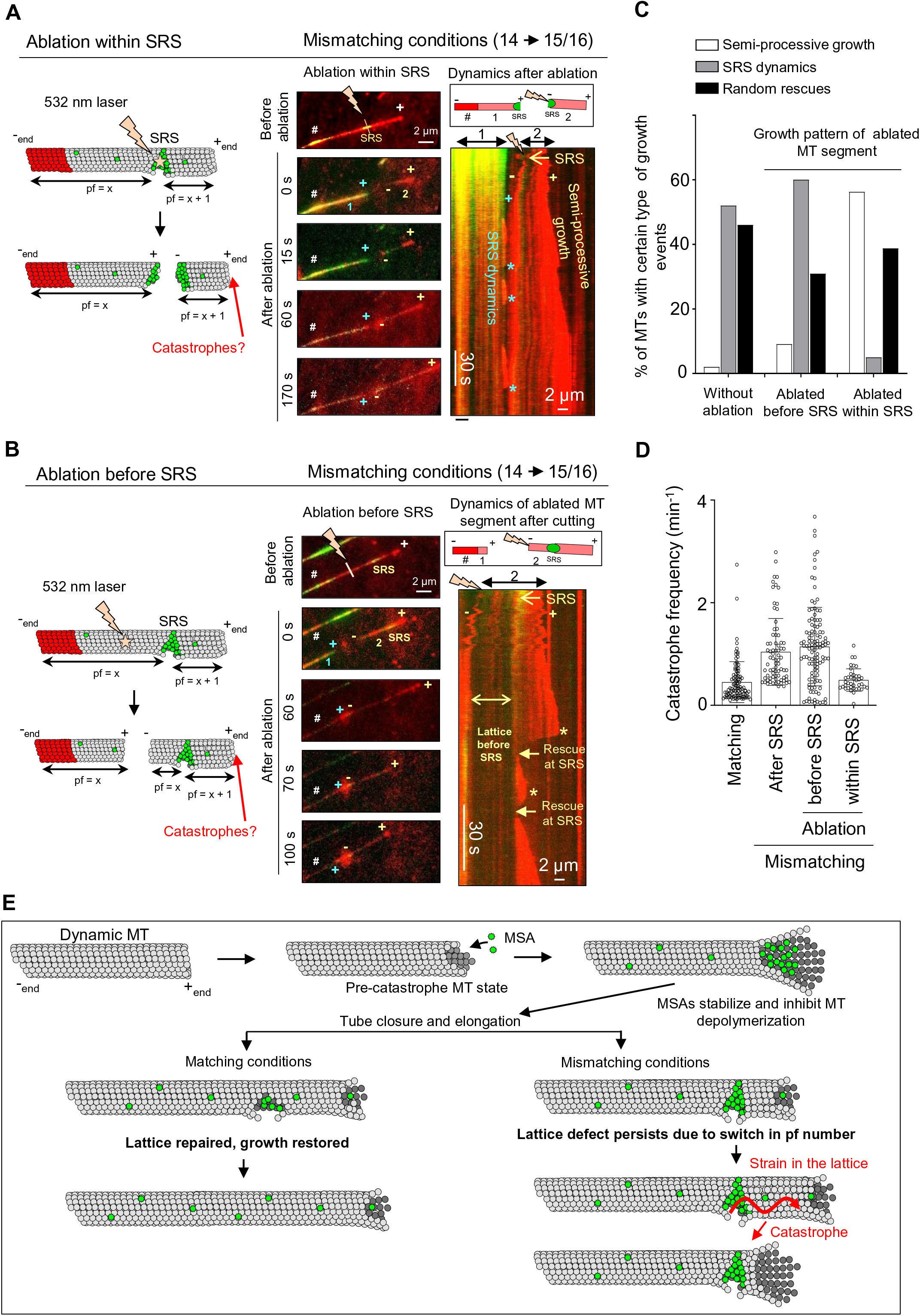
Microtubule severing at the stable rescue site suppresses catastrophes. **A, B**) Schematic representation and still images showing photoablation of microtubule regions within (A) or before (B) a drug accumulation zone (SRS) and kymographs showing microtubule dynamics of the severed parts. The assay was performed in the presence of GMPCPP seeds with 15 μM tubulin supplemented with 3% rhodamine-tubulin, 20 nM mCherry-EB3 and 100 nM Fchitax-3. The time-lapse images before photoablation show the presence of stable rescue sites; the site of ablation within (A) and before (B) the stable rescue site is indicated by a lightning bolt. The newly generated microtubule fragments (1 – seed-attached part, 2 – the part that is detached from the seed after ablation), microtubule plus (+) ends and the new plus (+) and minus (-) ends generated after ablation are indicated. In time-lapse images, # shows the position of the GMPCPP seed. In panel A, the kymograph illustrates the dynamics of both fragments (1 and 2, as shown in the scheme) generated after ablation. In panel B, the kymograph illustrates the dynamics of the severed microtubule fragment (2, as shown in the scheme). Asterisks highlight catastrophes, and rescues at the stable rescue site are indicated by arrows in panel B. The labels are in blue for fragment 1 (seed-attached microtubule part) and yellow for fragment 2 (the part detached after ablation). N = 5 independent experiments. **C)** Quantification of growth dynamics of the severed microtubule segment as represented in panels A and B. n = 60 and 55 for severing within and before a drug accumulation area, respectively. **D)** Catastrophe frequencies (calculated as frequency of all growth perturbations including “catching up” events) of microtubules growing in matching conditions (Fchitax-3 seeds + 100 nM Fchitax-3, n = 100), after the formation of stable rescue sites in mismatching conditions (GMPCPP seeds + 100 nM Fchitax-3, n= 76), microtubule fragments growing after photoablation in a region preceding the Fchitax-3 accumulation area (SRS) (n = 115) and microtubule fragments growing after photoablation within an Fchitax-3 accumulation area (SRS) (n = 39) in mismatching conditions. **E)** Model depicting microtubule lattice repair in matching and mismatching conditions. In pre-catastrophe state, MSAs stabilize depolymerizing protofilaments and inhibit depolymerization. In matching conditions, this leads to rapid repair of microtubule defects and restoration of microtubule growth. In mismatching conditions, lattice defects persist in spite of repair due to a switch in protofilament number. The presence of a defect might generate strain in the lattice which would affect the growing microtubule end and induce catastrophe.

When we severed microtubules directly at the site of Fchitax-3 accumulation (a stable rescue site), a part of the drug accumulation zone at the seed-attached microtubule fragment was often preserved after ablation. The plus end outgrowing from this zone displayed frequent catastrophes, typical for the SRS dynamics (white asterisks in Fig. 5A, Fig. S9A). However, the catastrophe frequency at the plus end of the newly generated microtubule fragment distal from the seed (with the minus end located at the former stable rescue site) significantly decreased and a large proportion of microtubules switched from SRS dynamics with long depolymerization episodes to semi-processive growth with short depolymerization episodes (Fig. 5A-D, Fig. S9A, Video S2).

In contrast, when we severed microtubules at a location preceding an Fchitax-3 accumulation zone, so that the stable rescue site with the flanking region on the minus-end side was preserved, microtubule plus ends distal from the severing site still underwent catastrophes, as was typical for the SRS dynamics (Fig. 5B-D, Figure S9B, Video S3). These data support the idea that drug-induced lattice discontinuities can exert an effect on the growth of microtubule plus end located at a significant distance away. To estimate this distance, we measured the average length of microtubule growth from the stable rescue site. This length was shorter than the average microtubule growth length after a seed, in line with the conclusion that catastrophe frequency after a stable rescue site is elevated (Fig. S9C). However, microtubules could still extend from a stable rescue site for 1-10 µm before undergoing a catastrophe (Fig. S9C). This indicates that conformational alterations or strain can propagate from a drug-induced defect to induce plus-end catastrophe at a distance encompassing hundreds of tubulin dimers.

## Discussion

In this study, we addressed how microtubule lattice defects induced by microtubule-stabilizing agents affect growth at the microtubule plus end. We made use of the fact that when microtubules are grown in the presence of non-saturating concentrations of MSAs, such as taxanes, the drugs strongly bind to microtubule ends in a pre-catastrophe state and thereby induce regions of increased microtubule stability, termed stable rescue sites (23). Importantly, these sites can contain “holes” in the microtubule wall, where fresh tubulin can be incorporated (23). The exact structural nature of these defects is likely to be complex. For example, recent cryo-EM work has shown that microtubules are not perfectly cylindrical but display strong local deviations from helical symmetry with different lateral contact geometries, and these deviations can be affected by taxanes (40). Local deviations from a cylindrical shape and additional types of microtubule lattice conformations, such as tubulin sheets (41) or protofilament flares (42) will likely have a major impact on the defect structure, stability and affinity for MSAs. Importantly, in the current study, we show that MSA-induced lattice defects represent areas of alterations in protofilament number, providing a simple geometrical reason for their persistence over time, despite continuous microtubule repair. This view is supported both by cryo-EM data and by measuring microtubule growth speeds, which we found to correlate with protofilament number.

The observation that an increase in microtubule protofilament number leads to a higher growth rate in the absence of a clear effect on the catastrophe frequency or depolymerization rate is unexpected and intriguing. In principle, one might expect that if the flux of tubulin subunits onto the end were independent of protofilament number, more protofilaments would lead to slower growth. Since we find that microtubules with more protofilaments grow faster, tubulin flux is probably not a major determinant of the microtubule growth rate in the conditions we use. Microtubules with higher protofilament number have a smaller angular separation between protofilaments, and this can affect the strength of lateral tubulin bonds and the ease with which they form. The current understanding of the molecular details of microtubule growth relies on combining experiments with modeling (43–48). For example, one recent model of microtubule polymerization assumes that growing microtubule tips have flared protofilaments, to the end of which tubulin dimers are added before the protofilaments associate with each other laterally (49). In this model, lateral interactions between tubulins were described by two parameters, an activation energy barrier and bond strength. Increasing protofilament number could reduce the lateral activation energy barrier and lead to stronger lateral bonds. According to this model, both effects would accelerate microtubule growth; however, these effects might potentially compensate each other during depolymerization, because strengthening of the lateral bonds would impede microtubule disassembly, whereas reducing the lateral activation energy barrier would promote it (49). Furthermore, microtubules with different protofilament number are expected to differ in the relative abundance of various types of lateral tubulin contacts (40), including “seam-like” contacts between α- and β-tubulin, and this will likely affect their polymerization kinetics.

Modification of protofilament number using MSAs allowed us to reproducibly generate lattice defects and explore the consequences they have on microtubule dynamics (Fig. 5E). Strikingly, we found that although a microtubule could polymerize beyond a persistent lattice defect for lengths up to several micrometers, microtubule plus end dynamics were affected, as the catastrophe frequency after the defect was significantly increased (Fig. 5E). A striking difference between microtubule plus ends grown from defect-bearing stable rescue sites and from stabilized seeds was that the latter could be efficiently “rescued” by an MSA once they entered a pre-catastrophe state. Drug accumulations leading to microtubule stabilization were ∼10-fold more frequent at the pre-catastrophe tips growing from seeds compared to microtubule ends entering a catastrophe after a preceding stable rescue site. These data indicate that some conformational aspects of a pre-catastrophe plus end during growth after a seed are different from those of a plus end elongating beyond a stable rescue site, and this might reflect differences in the catastrophe induction mechanism. However, our fluorescence-based measurements detected no significant change in microtubule end tapering and only a slight difference in the decay in EB3 intensity at pre-catastrophe ends of microtubules growing from seeds or after a stable rescue site. It would be interesting to use cryo-electron tomography to analyze the differences between growing microtubule ends in these conditions.

Strikingly, when microtubules were growing in matching conditions (when the MSAs used to stabilize the seeds and to elongate them had the same protofilament number preference), almost every growth perturbation resulted in rapid drug binding and tip repair, leading to semi-processive microtubule growth. These data show that the state and dynamics of the microtubule plus end depend not just on the conditions of polymerization, but also on the state of the preceding lattice (Fig. 5E). For example, microtubules growing in the presence of 100 nM Fchitax-3 from an Fchitax-3-stabilized seed and from a GMPCPP-stabilized seed after a stable rescue site encounter exactly the same concentrations of the drug and tubulin. Yet, in the first case, microtubule polymerization is somewhat faster and semi-processive, because although growth perturbations still occur just as in the absence of MSAs, microtubule ends transitioning to catastrophe are quickly repaired through drug binding and subsequent “catching up” events. In contrast, microtubules plus ends growing in the same conditions from a stable rescue site (i.e. after a lattice defect) undergo frequent catastrophes and are not repaired through MSA binding. Therefore, in mismatching conditions, catastrophes typically evolve into long depolymerization episodes.

Importantly, microtubule severing at the site of the defect made microtubules less catastrophe-prone, in line with the view that a lattice defect at the stable rescue site has propagating properties. The underlying mechanism is unclear, but it is possible that both tubulin extension or compaction in the axial direction (50, 51) and changes in angles between protofilaments due to heterogeneity of lateral contacts (40) might play a role. One surprising feature of the drug effects in our experiments is that they can be exerted at quite low drug binding densities (in some cases, less than 1 drug molecule per 8 nm of microtubule length or one tubulin “ring”). This suggests that effects of drug binding to a single tubulin dimer can propagate within the microtubule lattice. This is comparable to the reported effect of kinesin-1, where a few molecules binding to a microtubule could affect the structural properties of this microtubule (52). The finding that lattice defects have a propagating impact on microtubule plus end dynamics has important consequences for the concept of microtubule ageing. Previous work showed that the catastrophe frequency increases when a microtubule grows for a longer time, indicating that multiple steps are needed to induce a catastrophe (5, 6, 48). However, the nature of these steps is still unclear: they may be associated with a gradual change in the microtubule end structure (e.g. more tapering) (7, 9, 43), but may also occur within the lattice. Both types of changes might play a role, and in fact, our data suggest that the mechanistic basis of catastrophe induction may differ, as pre-catastrophe microtubule tips can be different both in terms of drug accumulation and EB3 decay. We found that the occurrence of drug-induced lattice defects associated with protofilament number mismatch leads to catastrophe. Whether protofilament number switching occurring in the absence of MSAs can also lead to catastrophe is currently unclear and deserves further investigation. Lattice defects including switches in protofilament number have been extensively documented in previous studies (10, 13–16). Since microtubule lattice defects can accumulate over time, they could potentially contribute to microtubule aging. Repair associated with microtubule defects has been described both in vitro and in cells, but until now, microtubule lattice discontinuities have been mostly linked to the formation of GTP islands, rescues and microtubule stabilization (18–21, 53). Here, we show that catastrophe induction is another important consequence of at least some types of lattice defects, and that lattice discontinuities can have a destabilizing effect on microtubules, a possibility that would need to be explored for defects occurring in the absence of MSAs in vitro and in cells.

It is also important to note that the drugs we employed, such as Taxol, Docetaxel and Epothilone B, are either used for therapies of cancer or considered for the treatment of neurodegenerative disorders (54–56). The concentrations of the drugs used in our assays are within therapeutically relevant range, and the knowledge that these drugs can differentially affect microtubule dynamics based on protofilament number preferences is relevant for optimizing their therapeutic applications.

## Materials and Methods

### Reagents and purified proteins

Taxol, 1,4-piperazinediethanesulfonic acid (PIPES), GTP, methylcellulose, glucose oxidase from *Aspergillus niger*, catalase from bovine liver, dithiothreitol (DTT), magnesium chloride, ethylene glycol-bis(2-aminoethylether)-N,N,N’,N’-tetraacetic acid (EGTA), potassium chloride, potassium hydroxide, κ-casein and glucose were obtained from Sigma-Aldrich. GMPCPP was obtained from Jena Biosciences. Biotinylated poly(l-lysine)-[g]-poly(ethylene glycol) (PLL-PEG-biotin) was obtained from Susos AG. NeutrAvidin was obtained from Invitrogen. Different types of labelled and unlabeled purified tubulin used in the assays were either purchased from Cytoskeleton or purified as described previously (57) for X-ray fiber diffraction experiments.

Docetaxel was procured from Sanofi-Aventis. Fchitax-3 was provided by Wei-Shuo Fang (State Key Laboratory of Bioactive Substances and Functions of Natural Medicines, Institute of Materia Medica, Beijing, China (58). Alexa_488_-Epothilone B was obtained from Simon Glauser and Karl-Heinz Altman (Department of Chemistry and Applied Biosciences, Institute of Pharmaceutical Sciences, ETH Zürich, Zurich, Switzerland (23). Discodermolide was synthesized as described previously (59).

### X-ray fiber diffraction

X-ray fiber diffraction images were collected in beamline BL11-NCD-SWEET of ALBA Synchrotron. Purified bovine tubulin (5 mg) was diluted to a final concentration of 100 µM in PM buffer (80 mM PIPES, 1 mM EGTA, 0.2 mM Tris, 1 mM DTT), containing 3 mM MgCl_2_, 2 mM GTP and 200 µM of the corresponding compound. Samples were incubated at 37 °C for 30 min to achieve maximum polymerization and then were mixed in a 1:1 volume ratio with pre-warmed PM buffer containing 3 mM MgCl_2_ and 2% Methylcellulose (MO512, Sigma-Aldrich). Final concentrations of protein, nucleotide and compounds were 50 µM tubulin, 1 mM GTP and 100 µM of compound. Samples were centrifuged for 10 s at 2000 g to eliminate air bubbles and transferred to a share-flow device (29, 60).

For each condition, 24 diffraction images were averaged and background subtracted using ImageJ software (version 1.51j8; Wayne Rasband, National Institutes of Health, Bethesda, USA). Angular image integrations were performed using XRTools software (obtained upon request from beamline BM26-DUBBLE of the European Synchrotron Radiation Facility (ESRF)). For average protofilament number determination, image analysis was performed as previously described (61) considering the absolute position of J_01_ for Taxol (0.0518 nm) as the reference for determining the relative peak positions for the remaining assayed conditions.

### Cryo-Electron Microscopy

*Microtubule polymerization:* For the analysis of protofilament number distribution, microtubules in the presence of GMPCPP or drugs were polymerized as described below in the section on microtubule dynamics, but in the absence of rhodamine- or biotin-labeled tubulin. For protofilament number transition analysis, microtubules were polymerized from GMPCPP seeds with 1 mM GTP, 15 μM tubulin, 20 nM mCherry-EB3 in the presence of 100 nM Fchitax-3 at 37℃ for 10 min.

#### Sample preparation

4 μl of each microtubule sample was applied to holey carbon grids (C-flat 2/2, Protochips) glow-discharged in air, before blotting and plunge-freezing using a Vitrobot Mark IV (Thermo Fisher Scientific) at 22 ℃ and 100% humidity.

#### Cryo-EM data collection

For protofilament number distribution analysis, cryo-EM micrographs of Taxol and GMPCPP microtubules were collected on a Tecnai T12 transmission electron microscope (Thermo Fisher Scientific) with a 4x4K CCD camera (Gatan), operating at 120 kV, image pixel size of 2.09 Å and defocus of around −5 μm. Cryo-EM micrographs of Fchitax-3 microtubules were collected on a G2 Polara transmission electron microscope (Thermo Fisher Scientific) with a K2 Summit detector, operating in counting mode and with a GIF Quantum LS Imaging Filter (Gatan) at 300kV, image pixel size of 1.39 Å and defocus range from −1.5 to −4 μm. 40 frames were motion corrected using MotionCor2 (62). For protofilament number transition analysis, cryo-EM images of microtubules were collected on a Tecnai T12 (Thermo Fisher Scientific) as above.

#### Cryo-EM data analysis

For protofilament number distribution analysis, protofilament number was determined by Moiré pattern visualization. 50-90 microtubules were selected for each dataset. For each microtubule population, the percentage of microtubules with a particular protofilament number was calculated.

For protofilament number transition analysis, microtubule diameters on each side of sheet-like lattice defects in Fchitax-3-microtubules were measured manually in Fiji. Diameters of equivalently separated sides of intact microtubules were measured as controls. To aid diameter measurement, Fourier transforms of microtubule regions were computed and filtered to enhance microtubule Moiré patterns. The filtered Fourier transforms were then inverse transformed to give filtered microtubule images.

### In vitro assay for microtubule dynamics

In vitro assay for microtubule growth dynamics was performed as described previously (35, 36). Briefly, as described earlier (23), microtubule seeds stabilized in the presence of GMPCPP (a slowly hydrolyzable GTP analog) were prepared by two rounds of polymerization and depolymerization in the presence of GMPCPP. A solution of 20 µM porcine brain tubulin mix containing 12% rhodamine-labeled tubulin/HiLyte 488^TM^ labeled tubulin and 18% biotin-labeled tubulin was polymerized in MRB80 buffer (80 mM K-PIPES, pH 6.8, 1 mM EGTA, 4 mM MgCl_2_) in the presence of GMPCPP (1 mM) at 37 °C for 30 min. After polymerization, the mixture was pelleted by centrifugation at 119,000 × g for 5 min in an Airfuge centrifuge. Obtained pellet was resuspended in MRB80 buffer and microtubules were depolymerized on ice for 20 min. The resuspended mixture was further polymerized in the presence of GMPCPP. After the second round of polymerization and pelleting, GMPCPP-stabilized microtubule seeds were stored in MRB80 containing 10% glycerol. For preparing microtubule seeds in the presence of microtubule stabilizing agents (MSAs), a solution of porcine brain tubulin (40 µM) mix containing biotin-labeled tubulin (18%) and rhodamine-labeled tubulin (12%) was polymerized in MRB80 buffer (80 mM K-PIPES, pH 6.8, 4 mM MgCl_2_, 1 mM EGTA) in the presence of GTP (1 mM) and indicated MSAs (20 μM) at 37 °C for 20 min. 20 μM MSAs (Taxol, Docetaxel, Alexa488-Epothilone B, Discodermolide and Fchitax-3) diluted in MRB80 were prewarmed to 37°C. Polymerizing tubulin solution was diluted 5 times with prewarmed 20 μM MSA solution and incubated further for 5 mins. The solution was centrifuged at 13200 × g for 15 min at 30 ⁰C. Obtained pellet was resuspended in 20 μM MSA solution (diluted in MRB80) and stored at room temperature with protection from light.

In vitro flow chambers were assembled on microscopic slides with plasma-cleaned glass coverslips using two strips of double-sided tape. Flow chambers were sequentially incubated with PLL-PEG-biotin (0.2 mg/ml) and NeutrAvidin (1 mg/ml) in MRB80 buffer. The chamber was further incubated with GMPCPP stabilized microtubule seeds followed by treatment with 1 mg/ml κ-casein in MRB80 buffer. The reaction mixtures containing 15 μM porcine brain tubulin supplemented with 3% rhodamine-tubulin when indicated, 20 nM mCherry-EB3 when indicated, 50 mM KCl, 0.1% methylcellulose, 0.2 mg/ml κ-casein, 1 mM guanosine triphosphate and oxygen scavenger mixture (50 mM glucose, 400 μg/ ml glucose oxidase, 200 μg/ml catalase, and 4 mM DTT in MRB80 buffer) with or without MSAs were added to the flow chambers after centrifugation in an Airfuge for 5 minutes at 119,000 × g. The chambers were sealed with vacuum grease, and microtubule dynamics was recorded using TIRF microscopy. All samples were imaged at 30 °C.

### Image acquisition by TIRF microscopy

Imaging was performed on a TIRF microscope setup (inverted research microscope Nikon Eclipse Ti-E) which was equipped with the perfect focus system (PFS) (Nikon) and Nikon CFI Apo TIRF 100x 1.49 N.A. oil objective (Nikon, Tokyo, Japan). The microscope was supplemented with TIRF-E motorized TIRF illuminator modified by Roper Scientific France/PICT-IBiSA Institut Curie, and a stage top incubator model INUBG2E-ZILCS (Tokai Hit) was used to regulate the temperature of the sample. Image acquisition was performed using either a Photometrics Evolve 512 EMCCD camera (Roper Scientific, final magnification 0.063 μm/pixel) or a Photometrics CoolSNAP HQ2 CCD camera (Roper Scientific, final magnification 0.063 μm/pixel) or a Prime BSI sCMOS camera (Teledyne Photometrics, final magnification 0.068 μm/pixel) and controlled with MetaMorph 7.7 software (Molecular Devices, CA). For simultaneous imaging of red and green fluorescence, we used a triple-band TIRF polychroic ZT405/488/561rpc (Chroma) and a triple-band laser emission filter ZET405/488/561m (Chroma), mounted in a metal cube (Chroma, 91032) together with an Optosplit III beamsplitter (Cairn Research Ltd, Faversham, UK) equipped with double emission filter cube configured with ET525/50m, ET630/75m and T585LPXR (Chroma). Images were captured with 5 frames/s in stream acquisition mode for Fig. S2B, 10 frames/s Fig. 5A, B, Fig. S9 A, B upper panel, 1 frame/s in time lapse mode for Fig. 3F and 1 frame/2s in time lapse mode for rest of the data.

### Analysis of microtubule dynamics in vitro

For Image analysis, ImageJ plugin KymoResliceWide v.0.4 (https://github.com/ekatrukha/KymoResliceWide (Katrukha, 2015)) was used for generating kymographs to represent the life history of microtubule dynamics. Microtubule dynamics parameters using kymographs were measured manually. The length and the duration of each growth event were measured as horizontal and vertical distances on the kymograph, respectively. Microtubule growth rate was determined as a ratio of these values. Catastrophe frequency was calculated as the inverse growth time. We note that for the microtubule dynamics data analyzed here, calculating catastrophe frequency by dividing the total number of catastrophes by the total time microtubule spent in growth produced numbers that were very similar. Depolymerization events (shrinkage of plus end/loss of EB3 intensity) with the length 0.2-0.5 μm and “catching up” events were considered as short growth perturbation events. Short growth perturbations including “catching up” events were considered as catastrophes when measuring catastrophe frequency. Depolymerization events shorter than 0.2 μm were not included in the analysis. We define dynamics as a stable rescue site (SRS dynamics) when a microtubule regrows at least 3 times from the same site after undergoing catastrophe. A random rescue is a single rescue event after a depolymerization episode that is longer than 0.5 μm. Two or more independent in vitro assays were performed for each reported experiment.

### Quantification of EB3 and Fchitax-3 intensity time traces

For the analysis, experiments were performed with 15 μM tubulin, 20 nM mCherry-EB3 and 100 nM Fchitax-3 in the presence of GMPCPP seeds (mismatch conditions) or Fchitax-3 seeds (matching conditions). This analysis was performed similarly to the procedure described in(23), apart from time trace alignment. Briefly, simultaneous two-color imaging of Fchitax-3/mCherry-EB3 was performed using an OptoSplit III beamsplitter (Cairn Research Ltd, UK) equipped with double emission filter cube projecting two channels on the camera chip simultaneously. Chromatic aberrations were corrected as described previously using calibration photomask(39). Registered videos were used to create kymographs by drawing segmented lines of 10-15 pixel width (0.65-1 μm) along episodes of drug accumulation on growing microtubules using KymoResliceWide plugin with maximum transverse intensity (http://fiji.sc/KymoResliceWide). On extracted kymographs, we outlined rectangular regions of interest (ROI) around observed accumulation event and exported both intensities to MATLAB. For each time point, we fitted mCherry-EB3 profile with sum of constant (lattice binding) and exponential decay functions (comet) convoluted with microscope’s point spread function (PSF):

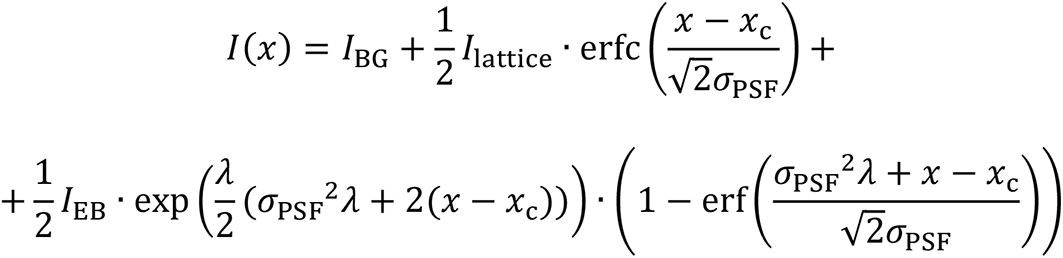

where fitting parameter *I*_BG_ corresponds to the intensity of background, *I*_lattice_ to the amplitude of the fluorescence intensity fraction associated with the lattice binding, *I*_EB_ to the amplitude of convolved exponential decay, *x*_c_ to the position of the maximum number of molecules in the molecules distribution (start of exponential decay position), *σ*_PSF_ to the standard deviation of microscope’s PSF and *λ* to the exponential decay constant. From the fitted function at each time frame we obtained maximum fluorescent intensity *I*_EB_MAX_(*t*).

Intensity of Fchitax-3 was fitted using Gaussian function of variable width with the addition of background. Total intensity was calculated as an integrated area under the fitted curve (without background intensity) and provided *I*_Fchitax3_(*t*). Both *I*_EB_MAX_(*t*) and *I*_Fchitax3_(*t*) intensity traces were normalized by the average of trace values above 80% percentile.

As a reference alignment, we chose Fchitax-3 channel for the time trace averaging, due to its clearly defined shape, and the EB3 time trace for each kymograph was shifted accordingly. Fchitax-3 traces were aligned using normalized cross-correlation. This pairwise function reports similarity between two temporal profiles depending on the displacement of one relative to another. It provides values in the range between zero and one, where higher values correspond to higher similarity. The global optimal alignment can be defined as a maximum of sum of cross-correlations between all profile pairs considering all possible displacements combinations. The number of combinations grows exponentially with the number of time traces and we found that time needed to explore the whole parameter space was too long for our datasets. Therefore, we devised an alternative algorithm searching for a local optimal alignment. We chose one time trace as a reference and registered all other time profiles to it, ensuring maximum of cross-correlation in each case. We repeated this procedure for all other time traces serving as a reference, and at the end chose the alignment with the maximum sum of cross-correlation functions among them. We observed that independent of the chosen reference profile, the majority of alignments provided very similar results with comparable cross-correlation score. This can be attributed to a distinctive sigmoid shape of Fchitax-3 temporal accumulation profile, making cross-correlation alignment converge to the same set of displacements. The corresponding MATLAB code performing the alignment and averaging, together with kymographs and ROIs used for the described analysis are available online (https://doi.org/10.6084/m9.figshare.c.5287663).

### Quantification of microtubule tip tapering and EB3 comet intensity

Experiments were performed with GMPCPP seeds in the presence of 15 μM tubulin supplemented with 6.7% HiLyte 488^TM^ labeled tubulin and 20 nM mCherry-EB3 (control, microtubule outgrowth from seeds) or the same mixture supplemented with 100 nM Discodermolide (mismatch conditions, microtubule outgrowth from stable rescue sites). Two-color imaging of HiLyte 488^TM^ labeled tubulin and mCherry-EB3, chromatic aberration correction of movies and kymographs generation were performed in the same way as described in the previous section. EB3 comet traces during episodes of growth ending with a catastrophe were manually outlined with polyline ROI, serving as initial estimation of a position. For each time point, intensity of HiLyte 488^TM^ labeled tubulin channel was fitted with the error function and background value:

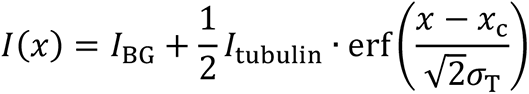

where fitting parameter *I*_BG_ corresponds to the intensity of background, *I*_tubulin_ to the amplitude of the fluorescent signal along microtubule, *x*_c_ to the position of the microtubule tip and *σ*_T_ to the degree of tip tapering convolved with microscope’s PSF. For the EB3 channel, maximum intensity of fitted comet *I*_EB_MAX_(*t*) was obtained as described in the previous paragraph. EB3 comet time trace was normalized by the average peak intensity over last 20 seconds before the catastrophe after smoothing it with 2.5s interval. The corresponding MATLAB code, kymographs and ROIs used for this analysis are available online (https://doi.org/10.6084/m9.figshare.c.5287663).

### Single molecule intensity analysis

Fchitax-3 molecules spontaneously binding to the coverslip during the reaction were used to measure single-molecule intensity (23). Briefly, two parallel flow chambers were made on the same coverslip. The first chamber was incubated with 100 nM Fchitax-3 without the reaction mixture. In second chamber, microtubule dynamics assay in the presence of Fchitax-3 stabilized microtubule seeds with 15 μM tubulin, 20 nM mCherry-EB3 and 100 nM Fchitax-3 was performed. For both the chambers, first 10-20 images of unexposed coverslip areas were acquired with the 100 ms exposure time using low laser intensity. In addition, a video of 300 frames exposing the same area with continuous laser illumination was recorded to induce photobleaching of Fchitax-3 molecules. Fluorescence intensities of Fchitax-3 molecules binding to the coverslip during reaction and in the solution of Fchitax-3 without the reaction mixture were detected and measured using ImageJ plugin DoM_Utrecht v.0.9.1 (https://github.com/ekatrukha/DoM_Utrecht). The fitted peak intensity values were used to build fluorescence intensity histograms. The intensities of Fchitax-3 molecules in the solution of Fchitax-3 without the reaction mixture and Fchitax-3 molecules transiently immobilized on the same coverslip during the reaction had the same intensity and showed the same single-step photobleaching behavior indicating Fchitax-3 molecules transiently immobilized to the coverslip during the reaction are single molecules. Further, integrated fluorescence intensities of Fchitax-3 molecules at the sites of drug accumulation during catch-up events in matching conditions were measured using ImageJ and were compared to Fchitax-3 single molecule intensity. These measurements were used to calculate the number of Fchitax-3 molecules at the drug accumulation site. The number of Fchitax-3 molecules per 8 nm of microtubule length was calculated considering Fchitax-3-microtubules as 15pf.

### Laser ablation of microtubules in vitro

Photoablation assays were performed on the TIRF microscope which was equipped with an ILas system (Roper Scientific France/PICT-IBiSA) and a 532 nm Q-switched pulsed laser (Teem Photonics). In vitro microtubule dynamics assay was performed in the presence of GMPCPP stabilized microtubule seeds with 15 µM tubulin supplemented with 3% rhodamine-tubulin, 20 nM mCherry-EB3 without (control) or with 100 nM Fchitax-3. Control microtubules were severed randomly by 532 nm focused laser beam. In mismatching conditions (GMPCPP seeds extended in presence of 100 nM Fchitax-3), microtubule lattices were ablated at the site of Fchitax-3 accumulations or in the region between the seed and an Fchitax-3 accumulation. Ablated microtubule parts display diffusive movements, but due to the presence of methylcellulose in the assays they stay close to the surface and could still be observed by TIRF microscopy. Using ImageJ software, maximum projections were drawn, which provide the life history of the ablated microtubule fragment. Kymographs were drawn using ImageJ plugin KymoResliceWide v.0.4.

## Statistical analysis

GraphPad Prism 7 was used to plot all the histograms and statistical analysis was done using non-parametric Mann-Whitney U-test. For Fig. 4B, two-sided Fisher’s exact test was performed.

## Data availability

All data that support the conclusions are available in the manuscript and/or available from the authors on request.

## Code availability

All MATLAB code, kymographs and ROIs used for the analysis are available online (https://doi.org/10.6084/m9.figshare.c.5287663).

## Acknowledgements

We thank G. F. Díaz for calf brains supply and staff of beamline BL11-NCD-SWEET (ALBA, Cerdanyola del Vallès, Spain) for their support with X-ray fiber diffraction experiment. We thank S. Kamimura (Chuo University, Tokyo, Japan) for kindly providing the share-flow device employed for fiber diffraction experiments. We thank W. S. Fang (Institute of Materia Medica, Beijing, China) for kindly providing Fchitax-3, S. Glauser and K.-H. Altmann (Institute of Pharmaceutical Sciences, ETH Zürich, Switzerland) for kindly providing Alexa_488_-Epothilone B and N. Gudimchuk (Lomonosov Moscow State University, Moscow, Russia) for helpful discussions. This work was supported by the European Research Council Synergy grant 609822 and the Netherlands Organization for Scientific Research (NWO) CW ECHO grant 711.015.005 to A.A., by a Biotechnology and Biological Sciences Research Council (BBSRC, BB/N018176/1) grant to C.A.M., and by grants “Bases moleculares de la regulación de microtúbulos y sus implicaciones en la neurotoxicidad producida por fármacos” from Fundación TATIANA, 20202020E301 from CSIC and PID2019-104545RB-I00 from Ministerio de Ciencia, Innovación y Universidades to J.F.D.

## Author Contributions

A.R. designed and performed experiments, analyzed data and wrote the paper. T.L. and C.A.M designed and performed cryo-EM experiments and analyzed the data; E.A. analyzed data; J.E.G. and J.F.D. performed X-ray fiber diffraction experiments; I.P. provided synthetic discodermolide; L.C.K contributed to the design of the experiments and analysis of the data and models; A.A. designed experiments, coordinated the project and wrote the paper.

## Legends to Supplementary Figures

**Fig. S1.**
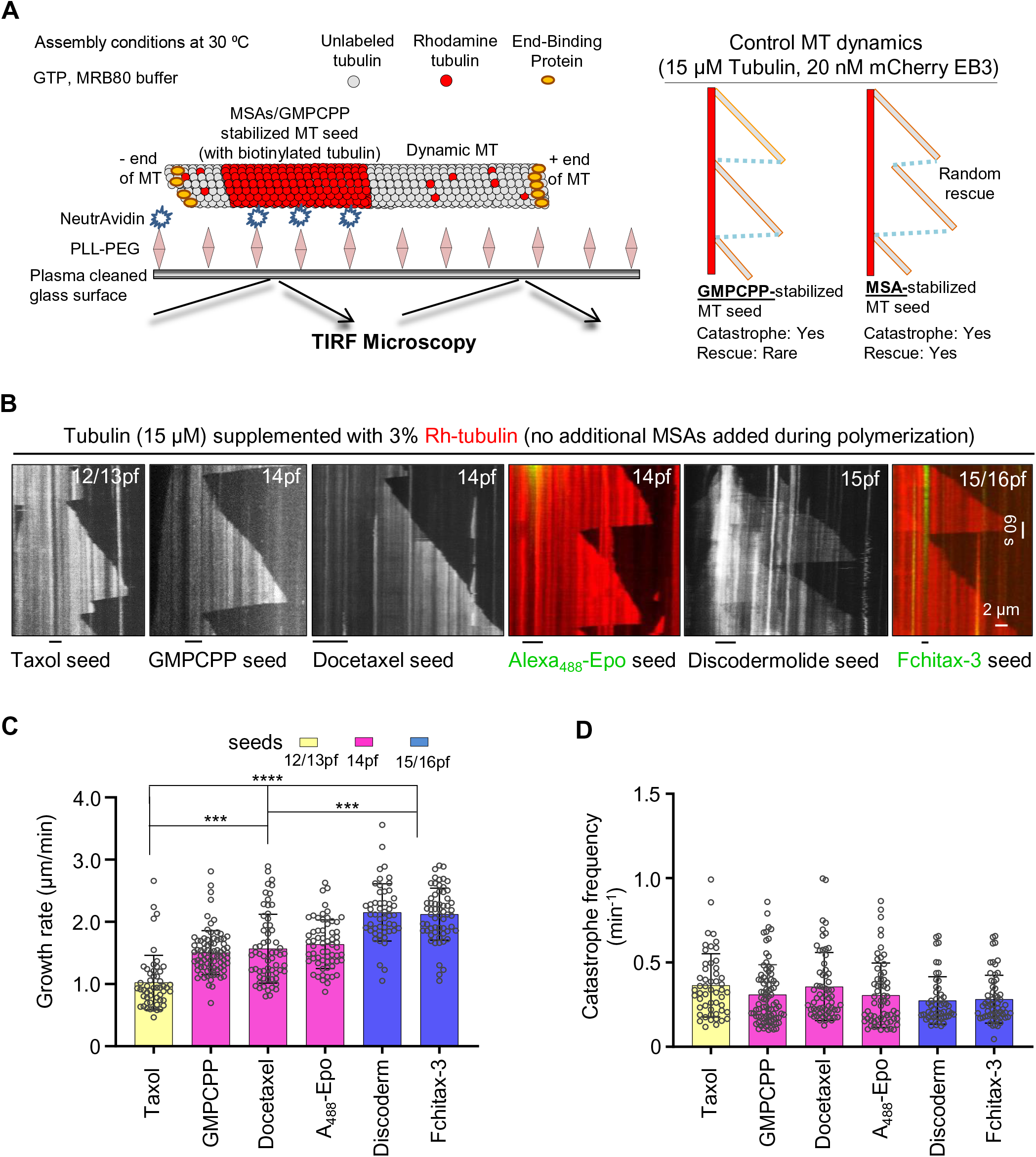
Effects of microtubule protofilament number on microtubule growth rate. **A)** Schematic representation of the in vitro microtubule dynamics assay and cartoons of kymographs illustrating microtubule growth events from a microtubule seed stabilized with GMPCPP or microtubule stabilizing agents (MSAs). **B)** Representative kymographs showing microtubule dynamics in the presence of seeds (stabilized with GMPCPP or the indicated drugs) with different protofilament numbers in the presence of tubulin (15 μM supplemented with 3% of rhodamine tubulin) in the absence of EB3 and without additional MSAs added during the reaction. **C, D)** Quantification of growth rates (C) and catastrophe frequencies (D) in the presence of seeds with different protofilament numbers as presented in panel B. From left to right, n = 52, 82, 62, 62, 51, 60 growth events. N = 3 independent experiments for Taxol, GMPCPP and Fchitax-3-microtubule seeds, N = 2 independent experiments for Docetaxel and Alexa488-Epothilone B-microtubule seeds, error bars represent SD; ***, p <0.001, ****, p <0.0001, Mann-Whitney U test.

**Fig. S2.**
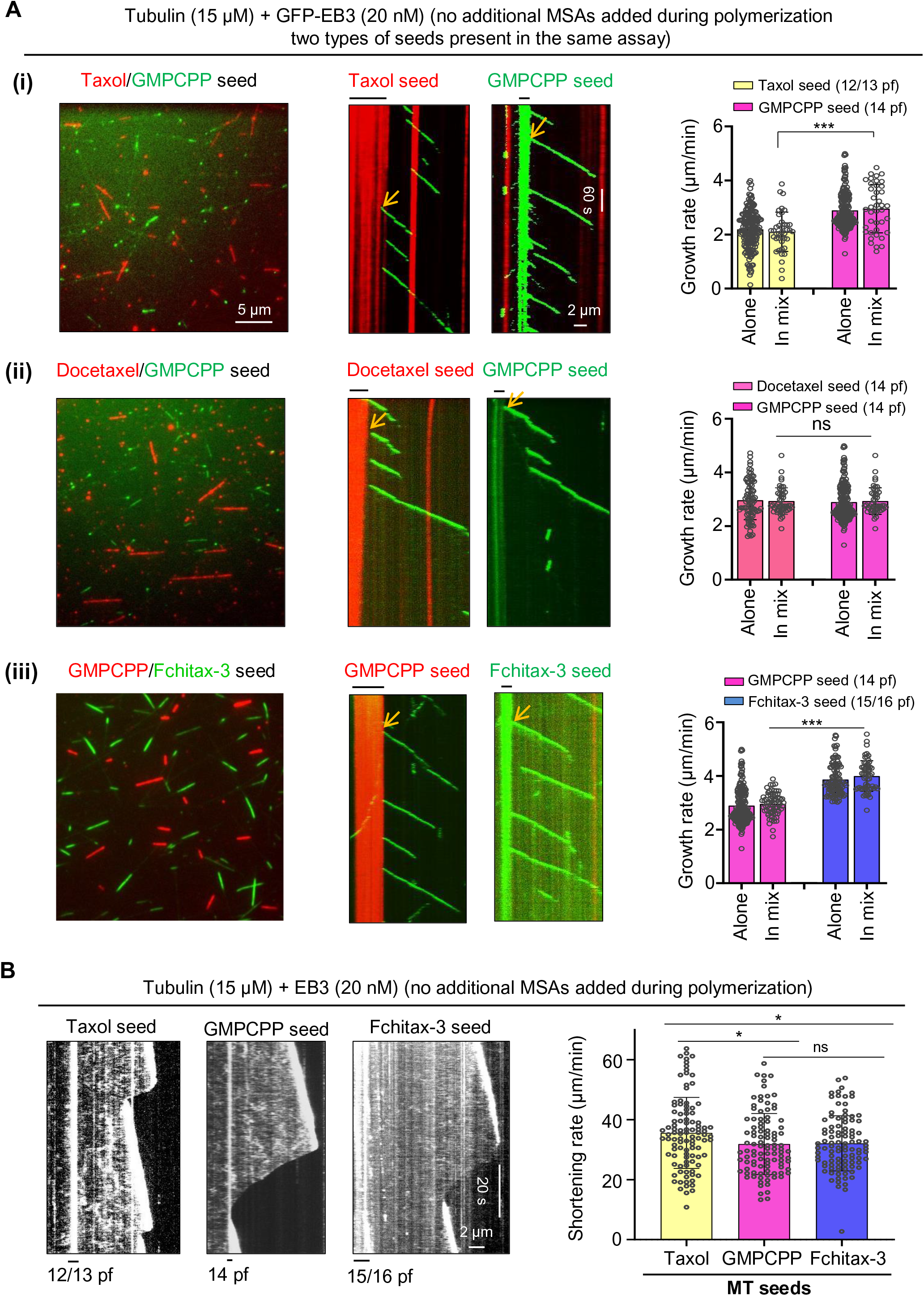
Effects of microtubule protofilament number on microtubule growth and depolymerization rates. **A)** Still images and representative kymographs showing the presence of two different kinds of seeds and microtubules growing from these seeds in the same assay. In the upper panel, Taxol (12/13pf) and GMPCPP (14pf) seeds were mixed (i). In the middle panel, Docetaxel seeds (14pf) were mixed with GMPCPP seeds (14pf) (ii). In the bottom panel GMPCPP seeds (14pf) were mixed with Fchitax-3 seeds (15/16pf) (iii). Respective bar graphs show the quantification of microtubule growth rate from different seeds during seed mixing conditions. Only growth events originating from seeds (highlighted by yellow arrows) and not from rescue sites were included in the quantification. For better comparison, data from Figure 1D (“alone”) are included in the present panels. From left to right-n = 193, 40, 196, 40 (panel i), n = 82, 47, 196, 47 (panel ii), n = 196, 66, 104, 60 (panel iii). **B)** Representative kymographs showing microtubule depolymerization events as indicated. Bar graph shows the quantification of shortening rates. N = 3 independent experiments, n = 100 for all the conditions, *, p <0.01, Mann-Whitney U test.

**Fig. S3.**
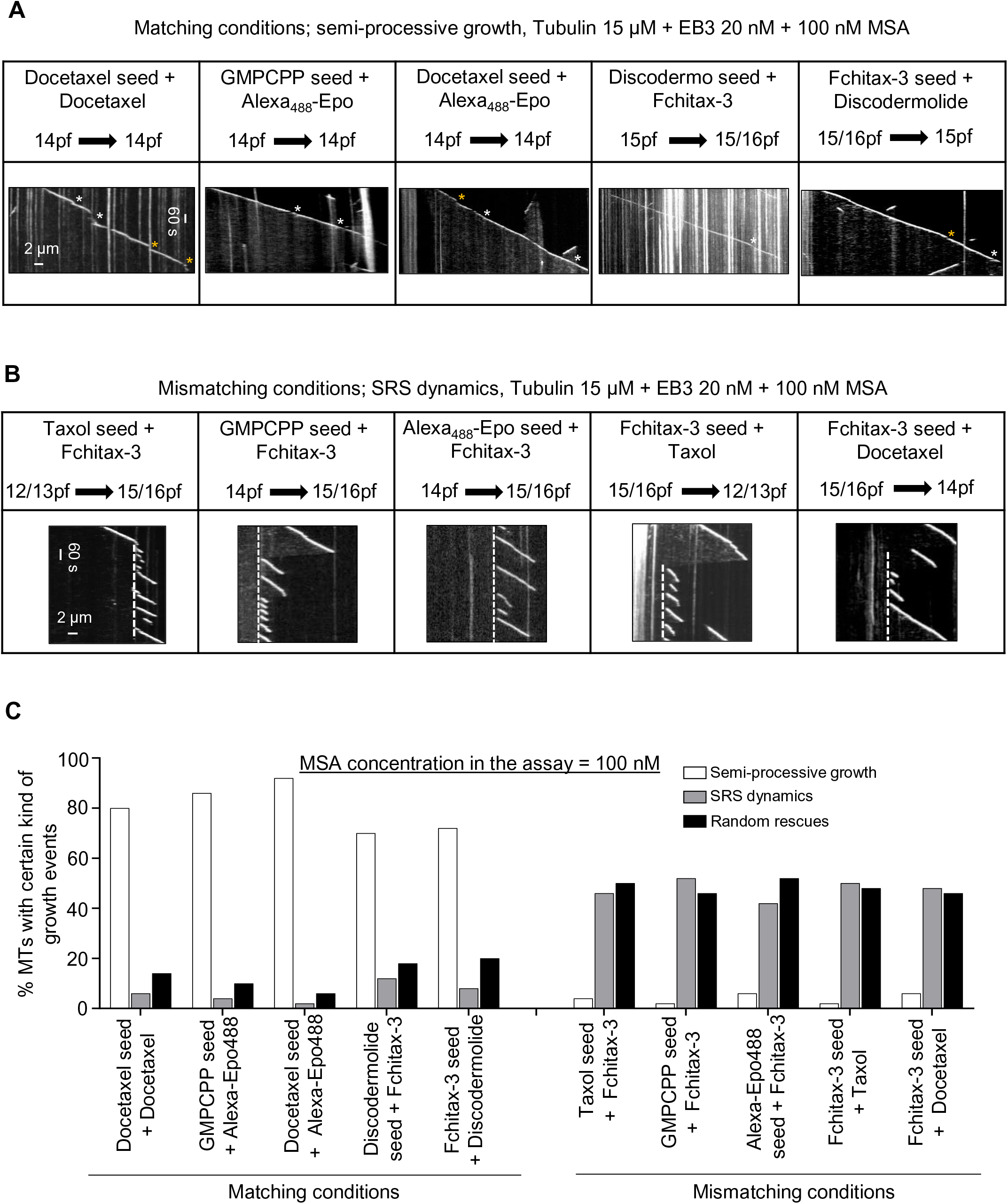
Effects of protofilament number mismatch between the seed and growth conditions on microtubule dynamics. **A, B)** Representative kymographs showing microtubule dynamics in the indicated conditions. N = 3 independent experiments. In matching conditions, short growth perturbation events followed by rapid rescues are highlighted (white asterisks highlight split comets and yellow asterisks highlight depolymerization events with the length 0.2-0.5 μm). In SRS dynamics, stable rescue sites in mismatching conditions are highlighted by white stippled lines. **C)** Quantification of microtubule growth patterns for conditions shown in panel A and B. n = 50 microtubule seeds in all conditions.

**Fig. S4.**
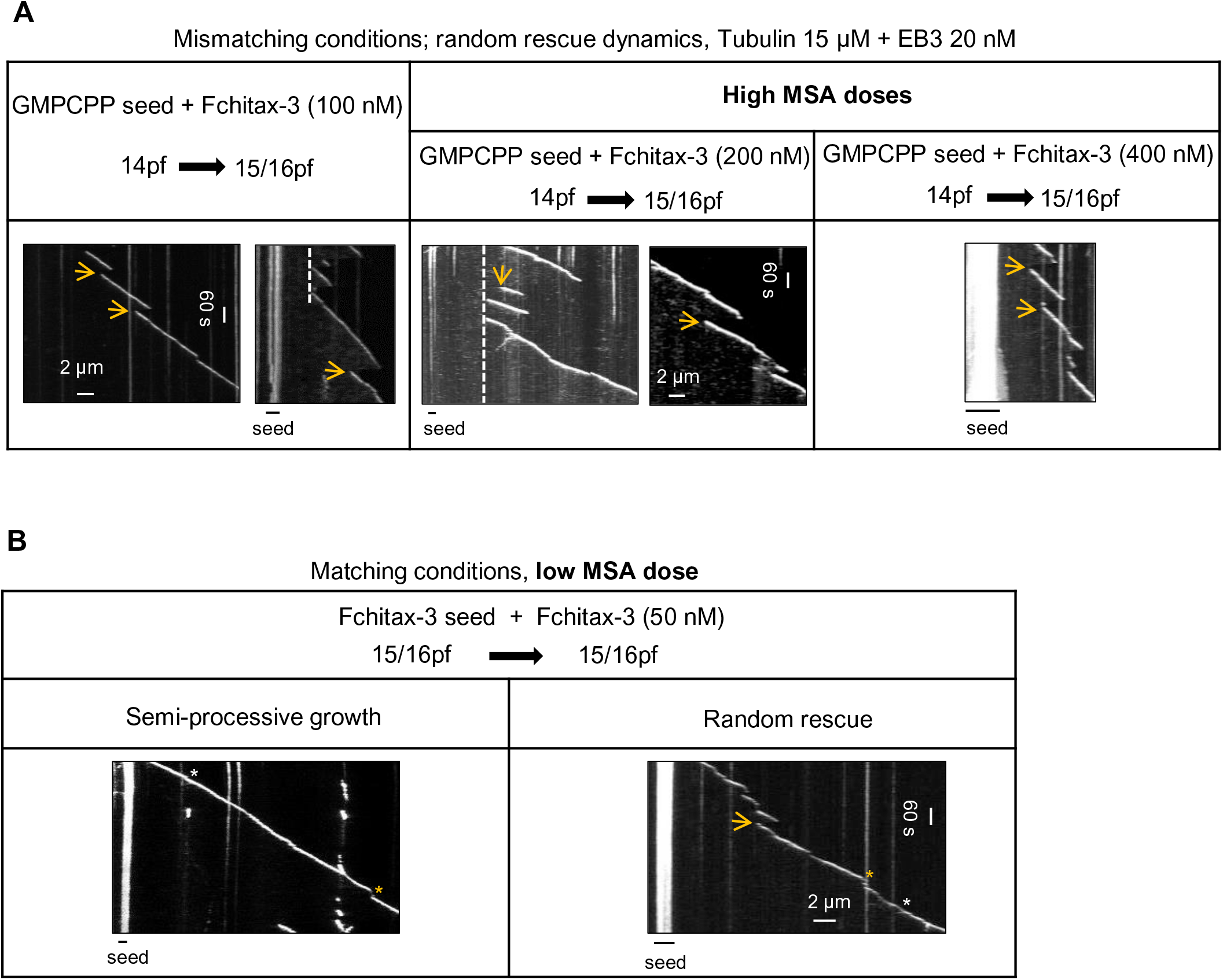
Effects of protofilament number mismatch between the seed and growth conditions on microtubule dynamics at different MSA concentrations. **A, B)** Representative kymographs illustrating microtubule dynamics in different conditions, as indicated. Random rescues are indicated with yellow arrows, white stripped lines highlight SRS, white asterisks highlight split comets and yellow asterisks highlight depolymerization events with the length 0.2-0.5 μm.

**Fig. S5.**
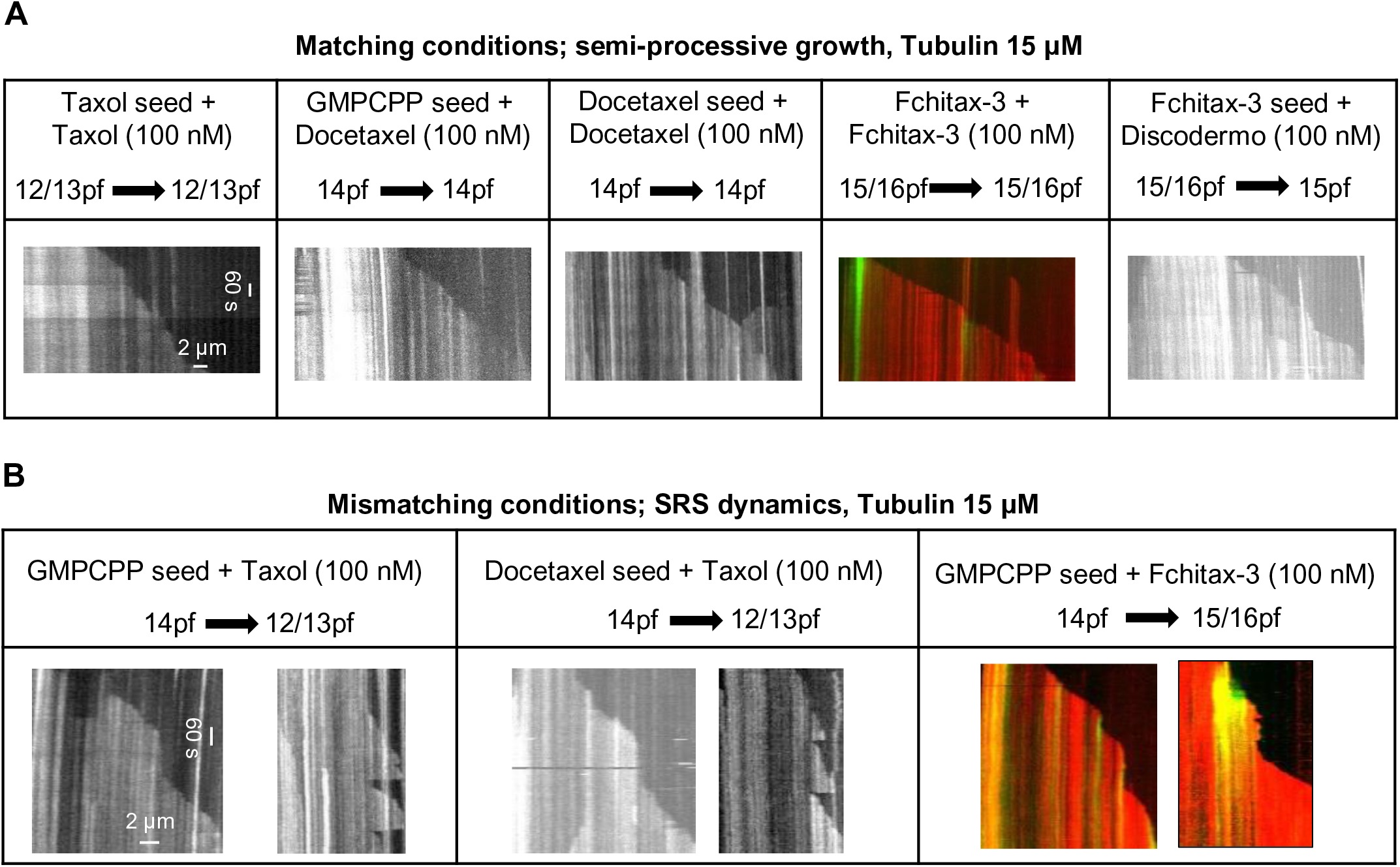
Effects of protofilament number mismatch between the seed and growth conditions on microtubule dynamics in the absence of EB3. **A, B)** Kymographs showing microtubule dynamics in the absence of EB3 for microtubules polymerized in the indicated conditions.

**Fig. S6.**
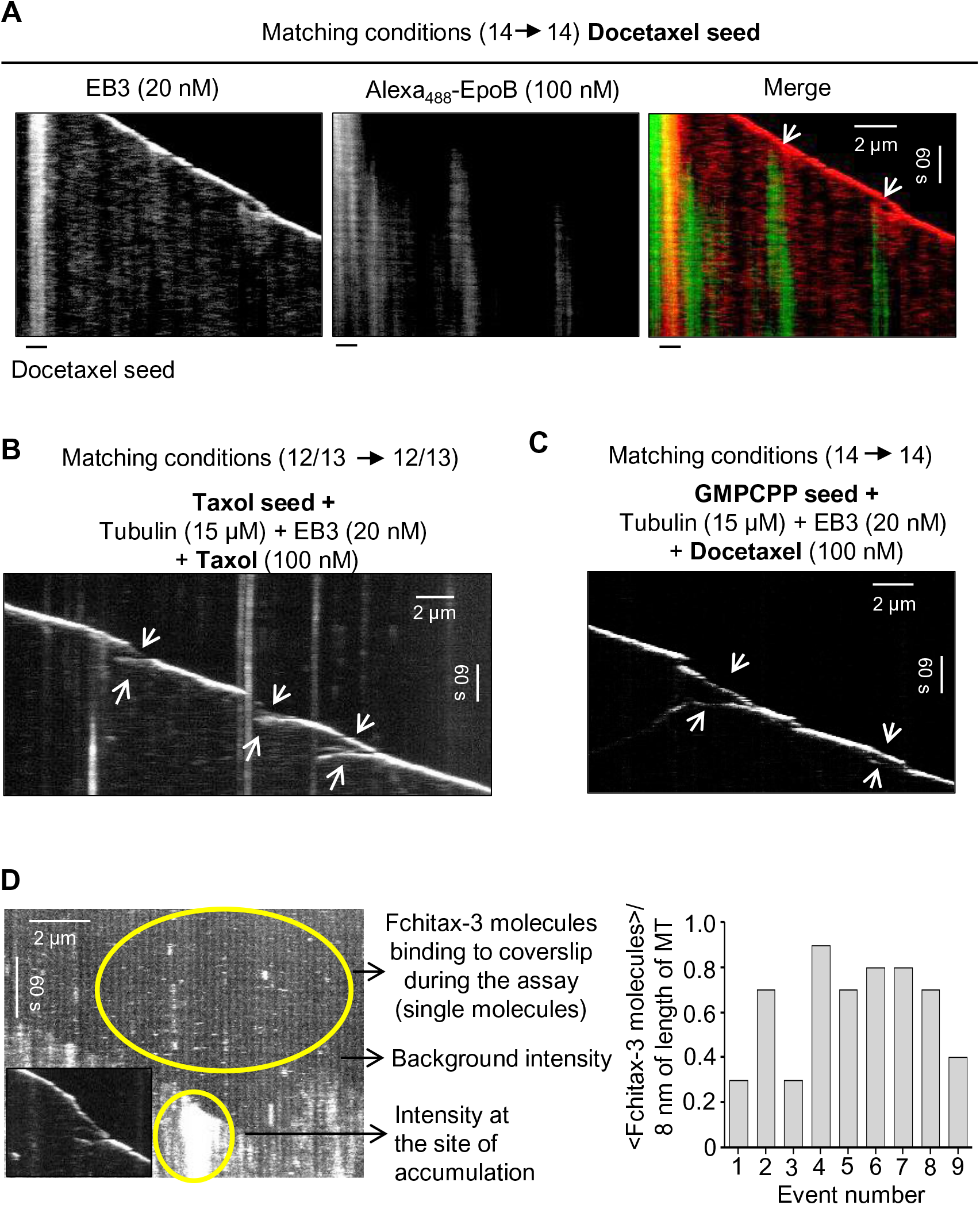
Characterization of microtubule growth perturbations in the presence of MSAs in matching conditions. **A)** Representative kymographs illustrating the dynamics of microtubules grown in matching conditions, from Docetaxel seeds in the presence of 15 μM tubulin and 20 nM mCherry-EB3 with Alexa_488_-Epothilone B (100 nM). White arrows indicate split comets. **B, C)** Representative kymographs showing split comets (“catching up” events, white arrows) in matching conditions as indicated. **D)** Kymograph illustrating the accumulation of Fchitax-3 molecules at the site of a split comet (inset) and free Fchitax-3 molecules binding to coverslip during the assay. Quantification of the number of Fchitax-3 molecules at the accumulation site is shown on the right.

**Fig. S7.**
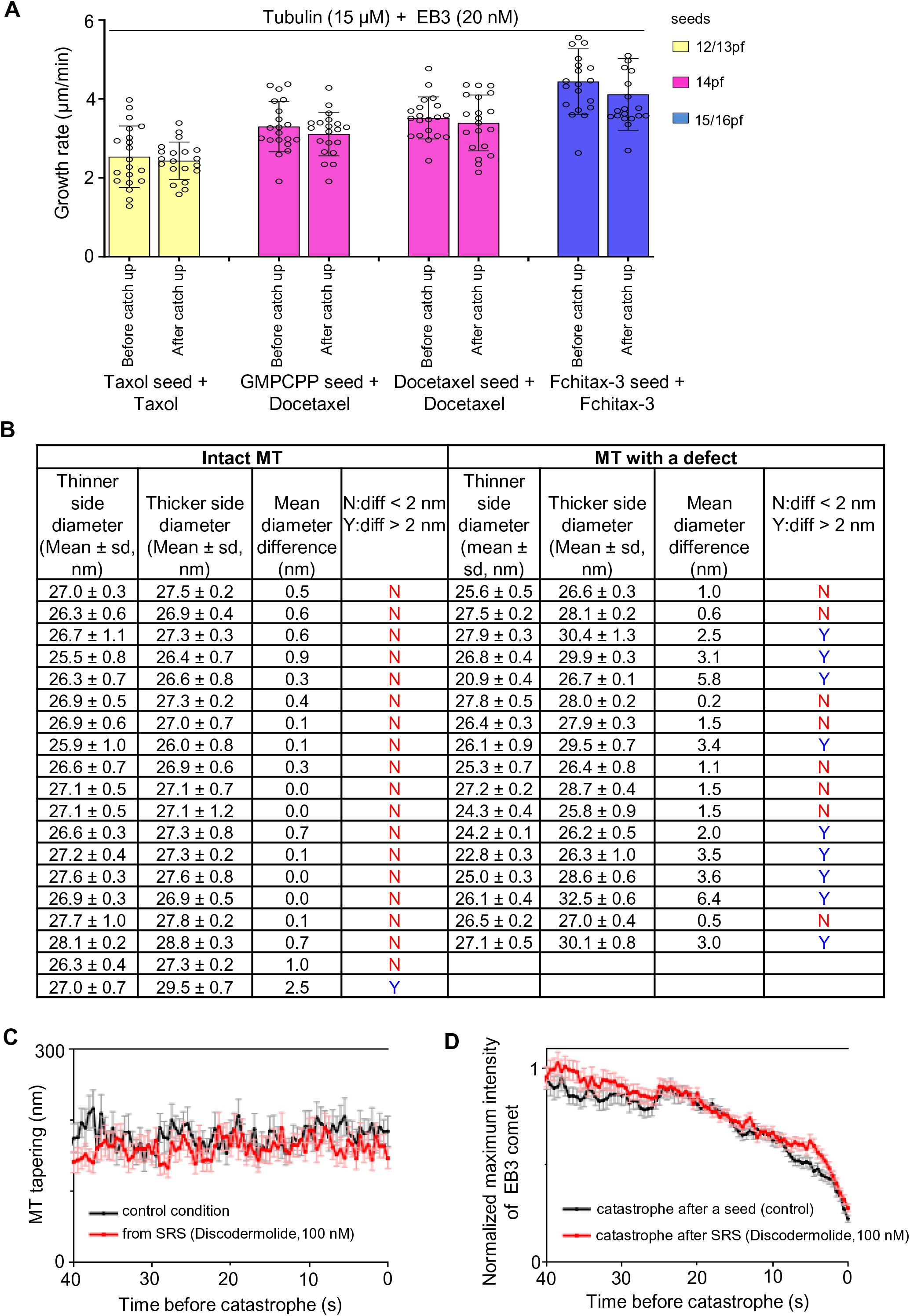
Analysis of changes in protofilament number associated with split comets and lattice defects. **A)** Quantification of growth rates before and after a “catching up” event in matching conditions. 20 events were analyzed per condition. **B)** A table showing the comparison of microtubule diameter on either side of a defect in microtubules grown from GMPCPP seeds in the presence of Fchitax-3. Microtubules with diameter difference of ≥ 2 nm were identified as microtubules with protofilament number transition. All possible microtubules with defects were picked from the dataset, excluding those with ice contamination and in crowded areas of the micrographs; as a control, an equivalent number of microtubules without defects were picked at random, also excluding those with ice contamination and in crowded areas of the micrographs. We used Fisher’s exact test, which is appropriate for the N of our data, to assess the difference in protofilament transition frequency in these microtubules. **C, D)** The degree of microtubule tip tapering (C) (derived from the fit to error function, with higher values corresponding to higher tapering) and averaged normalized maximum intensity of fitted EB3 comet (D) aligned by the moment of catastrophe. Black curves correspond to microtubules growing from seeds in control conditions (15 μM tubulin supplemented with 6.7% green tubulin, 20 nM mCherry-EB3 in the presence of GMPCPP seeds, 73 kymographs from 5 experiments). Red curves correspond to growth events from stable rescue sites (15 μM tubulin supplemented with 6.7% green tubulin, 20 nM mCherry-EB3 and 100 nM Discodermolide in the presence of GMPCPP seed, 64 kymographs from 5 experiments). Error bars represent SEM.

**Figure S8.**
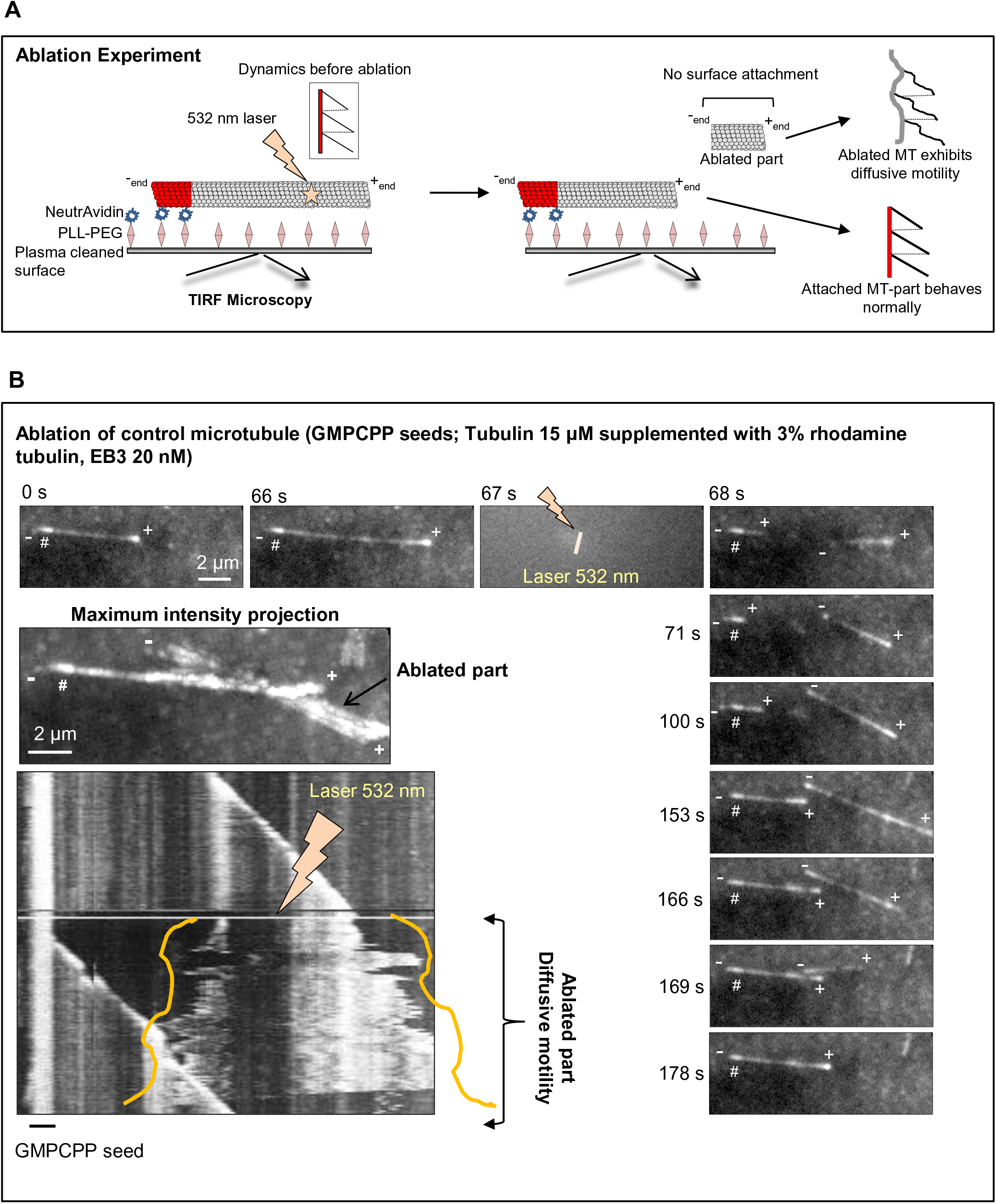
Laser mediated microtubule severing. **A)** A cartoon depicting laser (532 nm) ablation experiments with a TIRF-microscopy setup. The severed microtubule fragment is not attached to the coverslip surface and shows some diffusive motility. **B)** Still images showing photoablation of a control microtubule. The assay was performed in the presence of GMPCPP seeds with 15 μM tubulin, supplemented with 3% rhodamine-tubulin and 20 nM mCherry-EB3. # shows the position of the GMPCPP seed; the site of laser ablation is indicated by a lightning bolt. The plus and minus ends of the microtubule growing from the GMPCPP seed and the newly generated fragment are highlighted. The severed microtubule fragment is not attached to the coverslip surface and shows some diffusive motility, but stays in the same focal plane, as illustrated by a maximum intensity projection generated using ImageJ software. Kymographs were drawn using ImageJ plugin KymoResliceWide v.0.4.

**Figure S9.**
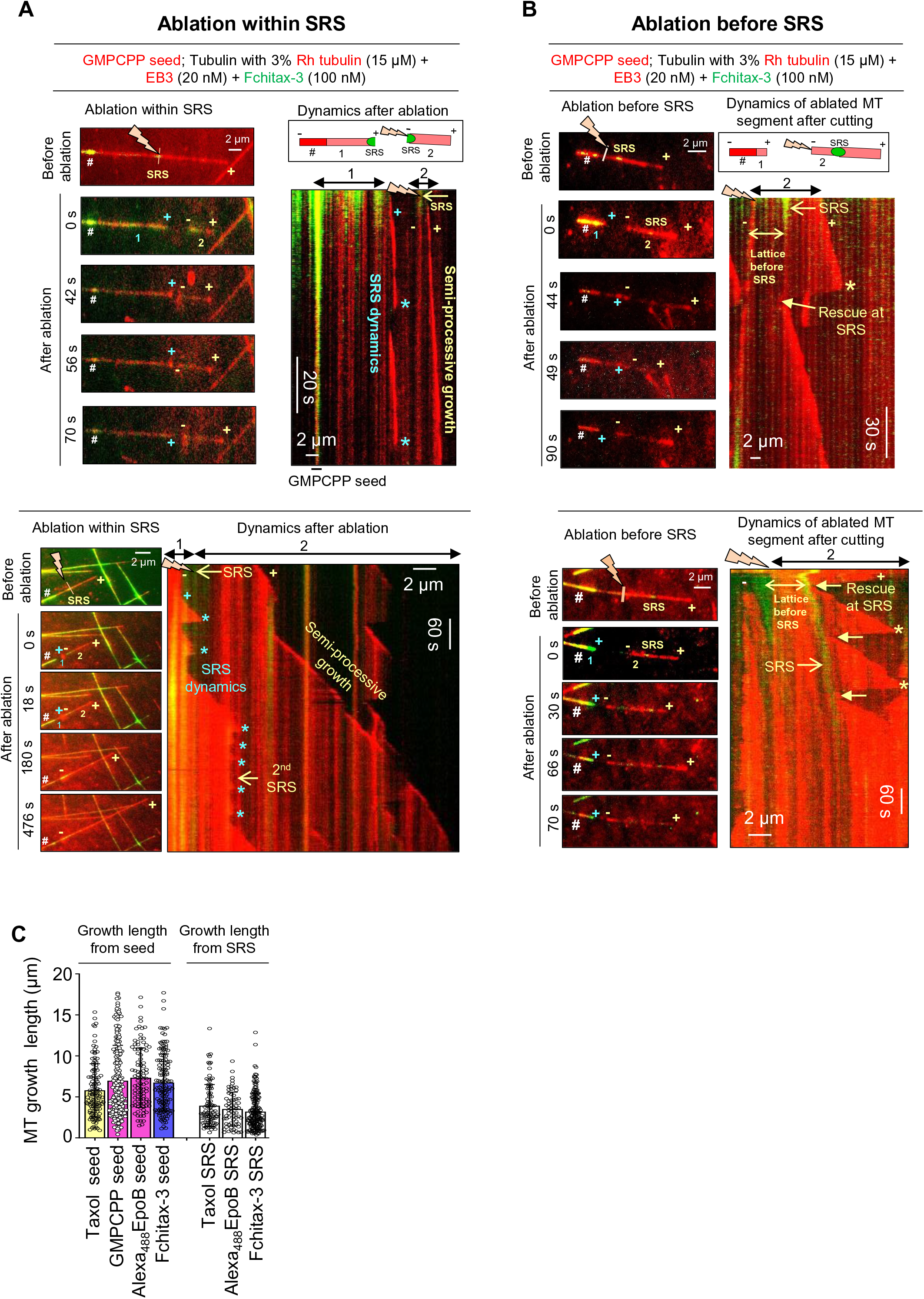
Microtubule severing within or before stable rescue site. **A, B)** Still images showing microtubule photoablation within (A) or before (B) a drug accumulation area (SRS) and kymographs illustrating microtubule dynamics of the severed microtubule segment after cutting. Upper panels, examples with faster acquisition (10 frames/s, stream acquisition). Lower panels, examples with slow acquisition (time-lapse acquisition with 1 frame/2s). The frame before ablation highlights the position of the GMPCPP seed (#), the plus (+) end of the microtubule, the position of Fchitax-3 accumulation within microtubule (SRS) and the site of laser ablation within or before SRS. After ablation, the severed microtubule parts (highlighted by 1 and 2), microtubule plus ends and the new ends generated after ablation are indicated. In panel A, kymographs illustrate the dynamics of both fragments (1 and 2) generated after ablation (as shown in the scheme). In panel B, kymographs illustrate the dynamics of the severed microtubule fragments (2, as shown in the scheme). Asterisks indicate catastrophes; rescues at the stable rescue site within the severed part are indicated by arrows in panel B. The labels are in blue for fragment 1 (seed-attached microtubule part) and yellow for fragment 2 (the part detached after ablation). **C)** Analysis of microtubule growth length for different conditions as indicated. From left to right: n = 118, 201, 96, 140, 92, 65 and 170.

## Legends to Supplementary Videos

**Video S1. Laser severing of control microtubules.**

The video illustrates photoablation of a control microtubule as depicted in Figure S8B. The experiment was performed in the presence of GMPCPP seeds with 15 µM tubulin supplemented with 3% rhodamine-tubulin and 20 nM mCherry-EB3. Laser ablation was performed using 532 nm laser on a TIRF setup. The video was acquired with a 1 s interval between frames and an exposure time of 100 ms. At the start, the positions of GMPCPP seed (#), plus (+) and minus (-) ends of the microtubule are shown. The moment and the site of ablation (!) are also marked at 67 s. The newly generated ends of the ablated part and the pre-existing ends are highlighted at 69 s. Scale bar, 2 µm. The video is representative of 5 independent experiments.

**Video S2. Laser severing of microtubule lattice within the Fchitax-3 accumulation zone (stable rescue site).**

The video illustrates photoablation of a microtubule within the Fchitax-3 accumulation zone (SRS), as depicted in Figure 5A. The experiment was performed in the presence of GMPCPP seeds with 15 µM tubulin supplemented with 3% rhodamine-tubulin, 20 nM mCherry-EB3 and 100 nM Fchitax-3. The video first shows a frame before photoablation, which highlights the position of the GMPCPP seed (#), the plus (+) end of the microtubule, the site of Fchitax-3 accumulation (SRS) and the site of laser ablation (!) within SRS. After photoablation within the SRS region, the video was recorded in a stream acquisition mode with an exposure time of 100 ms. After photoablation, only the red channel is presented. At the start of the image sequence after photoablation (0 s), the positions of the GMPCPP seed (#) and the microtubule ends are indicated. At 97.2 s, the growing ends of both parts are indicated. At 103.8 s, a catastrophe (*) of the seed-attached microtubule lattice is indicated. Scale bar, 2 µm. The video is representative of 5 independent experiments.

**Video S3. Laser severing of microtubule lattice before Fchitax-3 accumulation zone (stable rescue site).**

The video illustrates photoablation of a microtubule before the Fchitax-3 accumulation zone (SRS) as depicted in Figure 5B. The experiment was performed in the presence of GMPCPP seeds with 15 µM tubulin supplemented with 3% rhodamine-tubulin, 20 nM mCherry-EB3 and 100 nM Fchitax-3. The video first shows a frame before photoablation, which highlights the position of the GMPCPP seed (#), plus (+) end of the microtubule, the plus (+) end of the microtubule, the site of Fchitax-3 accumulation (SRS) and the site of laser ablation (!) before SRS. After photoablation, the video was recorded in a stream acquisition mode with an exposure time of 100 ms. After photoablation, only tubulin channel is presented. At the start of the image sequence after photoablation (0 s), the positions of the GMPCPP seed (#), and the microtubule ends are highlighted. After photoablation, both microtubule parts start growing (highlighted at 8.4 s and 66.4s). A catastrophe (*) of the ablated part is highlighted at 67.2 s. Scale bar, 2 µm. The video is representative of 5 independent experiments.

